# Coordination of spike timing among the neurons of the cerebellum

**DOI:** 10.64898/2025.12.03.692114

**Authors:** Mohammad Amin Fakharian, Elijah A. Taeckens, Alexander N. Vasserman, Alden M. Shoup, Reza Shadmehr

## Abstract

We think of neurons as either excitatory or inhibitory, but in the cerebellum, molecular layer interneurons (MLIs) inhibit Purkinje cells (P-cells) chemically, while exciting each other electrically. What is gained by electrical coupling of inhibitory neurons? Here we show that electrical coupling promotes a specific pattern of spike timing: as firing rates of two cells increase, the rate of spikes within 1ms of each other grows disproportionately while 2-4ms intervals are suppressed. If two MLIs produce spikes within 1ms of each other, the result is superposition of their individual effects on their target P-cell; a deep inhibition followed by a post-inhibitory rebound. This enhances the temporal precision in the P-cell’s next spike. However, when the MLIs spike 2-4ms apart, the two spikes interfere with each other, producing partial cancellation. Thus, electrical coupling of inhibitory neurons promotes production of spike intervals that enhance constructive superposition, reducing the variance of spike timing in the downstream neuron.

## Introduction

Neurons are classified as excitatory or inhibitory based on how they influence postsynaptic targets. However, in many regions of the brain, including the cerebral cortex ^1^, thalamus ^2^, and the cerebellum ^3,4^, inhibitory neurons excite each other via electrical coupling. In the cerebellum, Purkinje cells (P-cells) and molecular layer interneurons type 1 (MLI1s) are both inhibitory neurons that employ ephaptic coupling and gap-junctions ^4–6^. Does this electrical coupling play a role in control of behavior?

To answer this question, two problems must be solved. First, we must be able to simultaneously record from pairs of electrically coupled neurons that converge upon a single downstream neuron, a feat that is difficult because electrical connections exist among neurons that are at a distance of 100um or less ^5,7^. Second, because neurons modulate their firing rates during behavior, we must quantify coordination of spike timing ^8^, if in fact this coordination is present ^9^.

Here, we recorded from hundreds of P-cells and putative MLI1s (pMLI1s) in small neighborhoods termed “cliques” where inhibitory cell pairs of the same type excited each other ^10^. We discovered a mathematical pattern in the spike timing: as the firing rates of individual neurons increased during saccades, the rate of spikes that were no more than 1ms apart grew disproportionately with respect to chance, while the rate of spikes with a slightly longer interval (2-4ms) was suppressed or remained at chance.

To understand the purpose of this proclivity, we focused on triplets in which a pair of pMLI1s converged onto a target P-cell. When the spikes in the upstream pair were within 1ms of each other, they induced a deep suppression in the spiking probability of their target, then a post-inhibitory rebound. However, when the spikes were 2-4ms apart, they interfered with each other, producing partial cancellation. As a result, following a synchronous spike in the upstream pMLI1s the variance in the timing of the next spike in the downstream P-cell was reduced.

Whereas both P-cells and MLI1s employed electrical coupling, P-cells had an additional property: their spike timing was clock-like, remaining predictable across a wide range of firing rates ^11–14^. Simulations demonstrated that this regularity, in combination with ephaptic coupling, promoted P-cell synchrony at spike intervals of 1ms or less while suppressing asynchrony at 1-2ms intervals.

Thus, synchronous inhibition provided a means to control spike timing in the downstream neuron, suggesting that electrical coupling between P-cells provided the means for control of spike timing in the downstream nucleus neurons ^15^.

## Results

We trained marmosets to make saccades to visual targets (Fig. 1a) and recorded from 154 definitive P-cells (CS and SS) and 734 pMLI1s from lobules VI and VII of the vermis (Extended Data Fig. 1). We identified the P-cells based on their simple and complex spikes (SS and CS) and then used the distinct shapes of the CS waveforms in the dendritic tree and axon of each P-cell ^16,17^ to identify the molecular, Purkinje, and granular layers (Fig. 1c, Extended Data Figs. 1B & 2). In the molecular layer, we labeled neurons that inhibited the P-cells at 1ms latency or sooner as pMLI1s (Fig. 1e, Extended Data Figs. 3 & 4) ^18–20^. The pMLI1s exhibited a negative spike waveform in the molecular layer but a positive waveform in the granular layer, a feature of a pinceau (Figs. 1c, 1d, Extended Data Figs. 3 & 4) ^21^.

**Figure 1.**
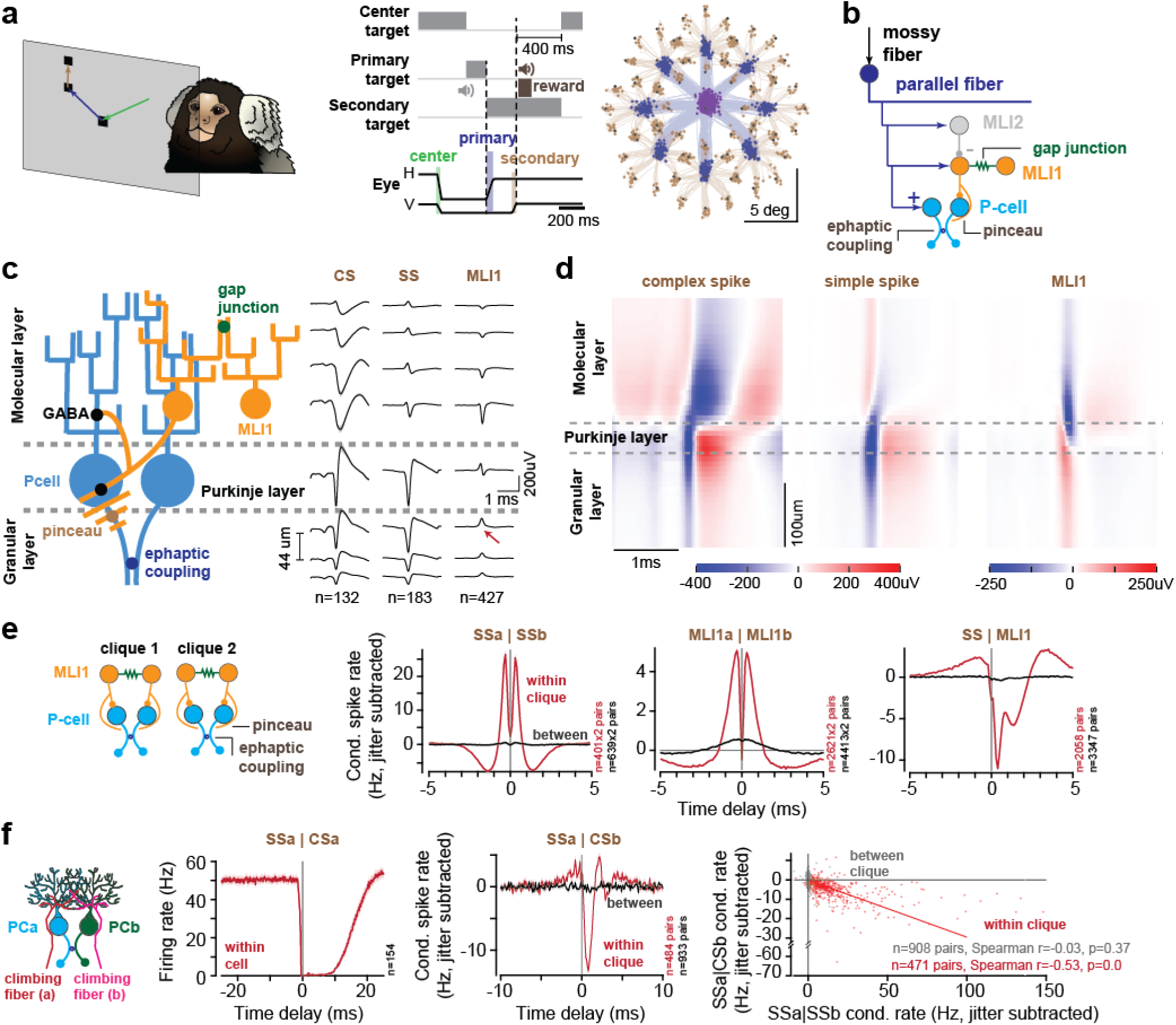
Spike interactions organized neurons into cliques within which neurons synchronized with each other. **(a)** Random corrective saccade task. **(b)** Network architecture, MLI2s inhibit MLI1s through GABAergic synapses, MLI1s inhibit P-cells via both GABAergic and electrical (pinceau) synapses; MLI1s are mutually connected through gap junctions; and P - cells excite each other through ephaptic coupling. **(c)** Schematic of neural organization and spatially aligned waveforms (projected perpendicular to the estimated P-cell layer, see methods & Extended Data Fig. 2). Spike waveforms for P-cells and pMLI1s are plotted at each corresponding layer. The data is from those subsets of cliques in which we had complex spikes recorded from P-cells at both the dendritic and axonal regions (see Extended Data Fig. 1B for all cliques). The upward pMLI1 spike near the P-cell layer (red arrow) is consistent with a pinceau. **(d)** Waveform alignment and layer identification. Average spatially aligned waveforms of complex spikes (left), simple spikes (middle), and pMLI1 spikes (right), obtained by identifying the Purkinje layer through the spatial boundary that separates dendritic and axonal components of complex spike waveforms within each clique (see methods). **(e)** Within and between clique jitter-subtracted conditional spike rates showing the likelihood that a spike occurred in one neuron given a spike in another neuron, across time delays. **(f)** Complex spike suppression of the same P-cell simple spikes (left), and ephaptic suppression of the neighboring P-cell simple spikes (middle). Ephaptic coupling of simple spikes and ephaptic suppression of complex spike on the simple spike are negatively correlated within pairs (two-sided spearman correlation test). Error bars are SEM.

In the molecular layer we identified pMLI2s (Extended Data Fig. 3, n=58) based on their inhibitory interactions with pMLI1s, and the fact that climbing fibers strongly and broadly excited the pMLI2s ^22,23^, but only weakly and briefly the pMLI1s ^20,24,25^. Unlike pMLI1s and P-cells, pMLI2s did not exhibit excitatory interactions with each other, and thus were not considered further.

### Spike interactions organized neurons of the cerebellar cortex into small networks

We next clustered the neurons into neighborhoods based on their interactions ^10^ (Extended Data Fig. 2). The spatial span of the probe allowed us to record from neurons in four Purkinje and molecular layers. However, only some of the cells interacted with each other. To visualize the interactions, we began with the jitter corrected ^26,27^ conditional probability of a cell producing a spike at time *t* + Δ, given that another cell produced a spike at time *t* (Fig. 1e). Using the results of this conditional spike rate (maximum absolute interaction with a delay of Δ ∈ {0,1,2,3} ms, 1ms time bin), we formed an adjacency matrix (Extended Data Fig. 2) where the numerical value in each element of the matrix was the strength of the spike interaction between the two cells. Next, we applied graph spectral clustering to the matrix and labeled cliques.

When two P-cells resided in the same clique, then an SS in one P-cell was followed by 25.4±1.19 Hz (mean±SEM) increase in the firing rate of another P-cell at 0.3ms latency, an interaction consistent with ephaptic coupling ^5^ (Fig. 1e, SSa|SSb, red trace). This interaction was an order of magnitude smaller when the two P-cells resided in different cliques (Fig. 1e, black trace, 0.56±0.06 Hz at 0.3ms).

When two pMLI1s were in the same clique, a spike in one pMLI1 was followed by 5.0±0.12 Hz increase in the firing rate of another pMLI1 at 0.3ms latency, an interaction consistent with gap junctions ^20^. These interactions were an order of magnitude smaller if the two pMLI1s were in separate cliques (Fig. 1e, MLIa|MLIb, 0.54±0.03 Hz at 0.3ms).

When a P-cell and a pMLI1 belonged to the same clique, the pMLI1 strongly inhibited the P-cell, producing a bimodal pattern of suppression: an initial inhibition in the SS rates at 0.4ms latency (-10.97±0.26Hz), followed by a second inhibition at 1.3ms latency (-5.76±0.21Hz, Fig. 1e, SS|MLI, also Extended Data Fig. 4), reproducing the pinceau and GABA-induced inhibitions in slice preparations ^21^.

The climbing fiber interactions with P-cells and pMLIs were also organized into cliques. CSs produced brief excitation of pMLI1s ^22^, and broad excitation of pMLI2s (Extended Data Fig. 3), likely via spillover ^18,19^ or direct synaptic contact ^23^, but only if they were in the same clique. CSs produced complete suppression of the SSs in the parent P-cell, but also briefly inhibited the neighboring P-cell (Fig. 1f), likely via ephaptic coupling ^17^. Notably, the larger the SS synchrony between a pair of P-cells, the greater the CS suppression from one P-cell upon the other (Fig. 1f, Spearman r=-0.53, p=0) ^17^. Thus, a P-cell that had strong SS ephaptic coupling with its neighboring P-cell also tended to have a climbing fiber that strongly suppressed the SS of that neighbor.

In summary, a spike in one P-cell was followed at 0.3ms delay by a 25Hz increase in spiking rate of another P-cell, but only if the two were in the same clique. A similar pattern of coordination was present among the pMLI1s.

### Spike intervals in electrically coupled cells were a linear function of firing rates

Recording sessions lasted 3.02±0.3 hours (median±MAD), with neurons modulating their firing rates during eye movements ^28^ and tongue movements ^29,30^. To test whether spike timing among cells was coordinated, we designed a new way to visualize the data, termed *joint-jitter plots* (Fig. 2c).

**Figure 2.**
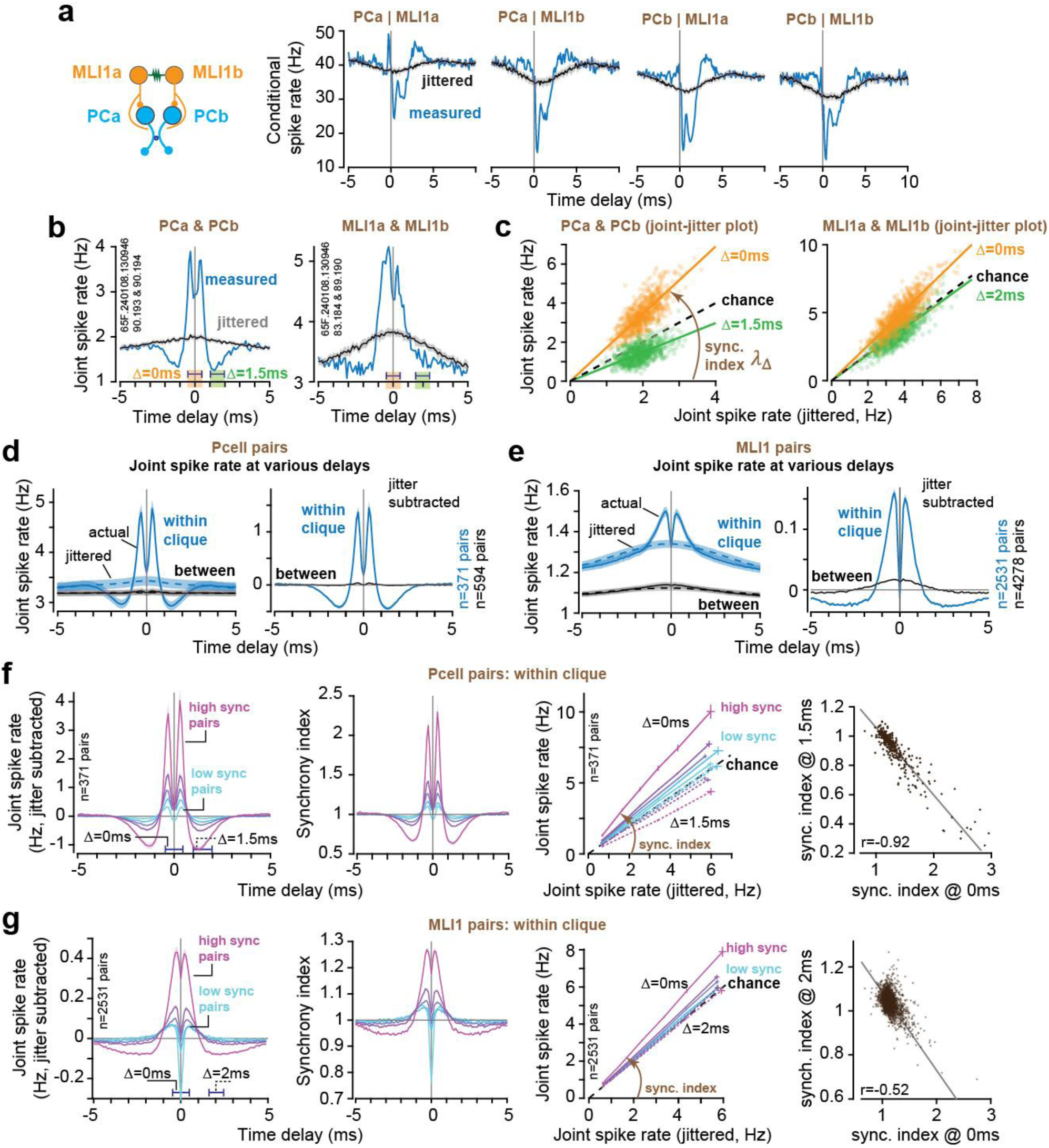
Spike timing among pairs of neurons exhibited a mathematical pattern. Data from the entire recording. **(a)** Conditional spike rate in a P-cell given a pMLI1 spike at 0ms (blue), compared to the same rate computed after jittering pMLI1 spikes (black; jitter window = 5ms, n=10 jitter iterations), shown for a sample P-cell and pMLI1 pairs in the third clique in the Neuropixels recording in Extended Data Fig. 2A, D. **(b)** Joint spike rate for the same P-cell pair (left) and pMLI1 pair (right), alongside their corresponding jittered controls (black). **(c)** Joint-jitter plots for the pairs in (b), each dot represents the number of synchronous spikes (within 1ms window; orange = 0ms, green = 1.5 & 2ms delay for P -cell and pMLI1 respectively) versus the average number of synchronous jittered spikes (n=10 jitte r iterations), calculated in non-overlapping 10s bins across the full recording. Lines indicate single parameter linear fit to the data points (slope is the synchrony index). **(d)** (left) Average raw (solid) and jittered (dashed) joint spike rate for all recorded P-cell pairs within (blue) and between (black) cliques. (Right) Jitter-corrected joint spike rate for the same groups. **(e)** Same as (d), but for all pMLI1 pairs. **(f)** From left to right: Jitter-subtracted joint spike rate of P-cell pairs grouped by connectivity strength. Synchrony index grouped by connectivity strength. Joint-jitter plots at 0ms (solid) and 1.5ms (dashed) delays for each group. Synchrony indices at 0ms vs. 1.5ms delay (each dot is one pair). **(g)** Same analyses as in (f), but for pMLI1 pairs and 2ms delay. Error bars are SEM.

Consider a pair of P-cells (PCa and PCb, Fig. 2a), and a pair of pMLI1s (MLIa and MLIb) that reside in the same clique. These two pMLI1s inhibited both P-cells, with the inhibition reaching its peak at a latency of less than 0.5ms after the pMLI1 spike, a pattern consistent with ephaptic inhibition via pinceau ^21^. To measure coordination of spike timing, we computed the joint probability of spiking in a window of size *w* = 0.1ms and then shifted this window across time delays Δ (Fig. 2b, blue line). As a control, we jittered the spikes in one of the P-cells and recomputed the joint probability (Fig. 2b, gray line). The joint probability for PCa & PCb exceeded the jittered values at ±0.4ms delay, then fell below those values at ±1.5ms delay (Fig. 2b, left subplot), a pattern consistent with ephaptic coupling between the two P-cells ^5^.

To visualize how the joint spike rate varied as a function of firing rates, we generated a new spike train that placed a spike only when both P-cells spiked during a 1ms window centered at Δ = 0ms delay. As a control, we jittered the spikes in one of the neurons, then computed the rate of the joint spiking in 10s windows of time and plotted it against the rate from the jittered data at the same window of time. We named this the *joint-jitter plot* (Fig. 2c). The x-axis presents the rate of joint spikes that are expected by chance. The y-axis is the rate that is observed in the data. For these two P-cells, at Δ = 0ms delay the joint rate exhibited a slope of greater than one (joint/jitter ratio, mean±SEM: 1.72±0.01, 1=chance), and at Δ = 1.5ms delay the joint rate had a slope of less than one (joint/jitter ratio, 0.75±0.007, Fig. 2c, left subplot). A similar pattern was present in the two pMLI1s. The rate of joint spiking at Δ = 0ms grew linearly as a function of the rate of the jittered data and had a slope greater than one (Fig. 2c, right subplot, joint/jitter ratio, 1.3±0.006). In contrast, at Δ = 2ms, the rate of synchronous spikes was near chance (joint/jitter ratio, 0.96±0.005).

Thus, there was a pattern in how the spikes of P-cell pairs, and MLI1 pairs, were organized in time: the probability of neuron *b* spiking at time *t*, and neuron *a* spiking at time *t* + Δ, was approximately a linear function of the jittered rates:

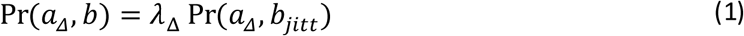

The term *λ*_Δ_ is termed synchrony index and is the slope of each line in Fig. 2c. Eq. (1) implied that as the firing rates of the two neurons changed, the rate of spikes that were Δ ms apart from each other remained a constant multiple of the individual firing rates (because Pr(*a*, *b_jitt_*) ≈ Pr(*a*) Pr(*b*)). For example, as the firing rates of two P-cells increased, so did the number of spikes at Δ = 0ms latency with respect to chance, but with a slope that was greater than one. That is, the rate of synchronous spikes grew faster than expected by chance.

Eq. (1) is noteworthy because its predictions differ from those of a recent paper. Herzfeld et al. ^8^ suggested that as P-cell firing rates changed, the difference between joint and jittered rates, termed covariance, remained constant. In contrast, Eq. (1) states that as firing rates changed, the ratio of the joint to the jittered probabilities remained constant, not covariance, consistent with in-vitro findings in P-cells ^5^.

Thus, if covariance is constant, we would expect the joint rates to rise in parallel to the jittered rates. If synchrony index is constant, we would expect the slope of the joint rate to remain constant as a function of jittered rates. It is also possible that neither of these predictions are correct.

We tested the predictions of Eq. (1) by computing the joint spike rate at various delays across the entire recording for all P-cell pairs (Fig. 2d, left plot, solid blue line), and pMLI1 pairs (Fig. 2e, left plot). The joint rate in P-cell pairs within a clique, with respect to chance, exhibited a sharp peak at Δ =±0.3ms delay (joint: 4.88±0.16Hz vs. jittered: 3.43±0.09Hz, Fig. 2d), then a minimum at Δ =±1.4ms delay (joint: 2.93±0.09Hz vs. jittered 3.38±0.09Hz). However, between cliques the joint rate was near chance (at 0.3ms joint: 3.23±0.06Hz vs. jittered 3.2±0.06Hz, and at 1.4ms joint: 3.18±0.06Hz vs. jittered 3.19±0.06Hz). The pMLI1s exhibited a similarly enhanced rate of joint spiking within cliques, and a small but non-zero rate of joint spiking between cliques (Fig. 2e, right).

Among the cells within a clique, some pairs exhibited much stronger joint spiking at Δ = 0ms than others. We divided the within clique pairs into five groups based on the strength of their synchrony index (Fig. 2f, left subplot), then made joint-jitter plots for each group. The joint-jitter plots revealed that as P-cell firing rates increased, the rate of joint spikes at Δ = 1.5ms remained below chance (Fig. 2f, third subplot). The greater the synchrony index at Δ = 0ms delay, the smaller the synchrony index at Δ = 1.5ms delay (corr: -0.92, p = 4.5×10^−149^ Fig. 2f, right subplot). In other words, the P-cells that had strong synchronous spiking also exhibited strong suppression of asynchronous spikes.

Synchrony index stratified pMLI1 pairs based on the strength of their coupling, while the coupling strength defined the slope of the joint-jitter relationship. Like the P-cells, the greater the synchrony index at Δ = 0ms delay, the smaller the synchrony index at Δ = 2ms delay (corr: -0.52, p = 7.74×10^−176^, Fig. 2g).

In summary, spike intervals in pairs of P-cells, and pairs of pMLI1s, exhibited a mathematical pattern: the probability that a pair of neurons would generate spikes at delay Δ from one another grew as a function of their jittered rates, with a slope that was greater than one for Δ = 0ms delay, but less than one for slightly longer delays (slope 1=chance). This means that as two neurons within a clique increased their firing rates, they promoted the spikes that were within 0.5ms of each other, while suppressing (in case of P-cells) or maintaining at chance (in case of pMLI1s) the spikes that were at 1.5ms of each other. The stronger the promotion at 0ms, the stronger the suppression at longer delays.

### Coordination of spike timing peaked near the onset of saccade deceleration

Fig. 3a presents the firing rates of P-cells and pMLI1s for saccades in various directions. Here, *θ* is the saccade direction for which complex spike activity was maximum ^28,31^. The direction *θ* is important because it specifies the direction of the potent vector of the clique ^10,32^. This means that when a P-cell is suppressed during a saccade, the eyes are pulled in direction *θ* ^33^. The P-cells as a population showed an early increase in their firing rates for saccades in direction *θ* + 180, and a late increase for saccades in direction *θ*. In contrast, the pMLI1s increased their rates for all saccades, displaying a slight shift in timing as a function of direction.

As P-cell firing rates changed during saccades, so did the rate of joint spikes (within 0 ± 0.5ms of each other, Fig. 3b, left column). However, regardless of saccade direction, the joint rate remained constrained to a line as a function of the jittered rates (Fig. 3b, right column, Extended Data Fig. 5). This is the static part of the pattern, which reiterates Eq. (1). The dynamic part of the pattern was unmasked when we subtracted the jittered rate from the joint rate (i.e., synchronous rate above chance). Because the slope in the joint-jitter function was greater than one, the rate of synchronous spikes above chance increased as the firing rates increased (Fig. 3c, left column). For the P-cells, the rate of synchronous spikes reached a peak before deceleration onset for saccades in direction *θ* + 180, i.e., the direction opposite to the potent vector. For the pMLI1s, the synchronous rate reached a peak just before deceleration for all saccade directions. Crucially, as predicted by Eq. (1), synchrony index *λ*_Δ_ remained roughly constant during saccades (Fig. 3c, right column).

**Figure 3.**
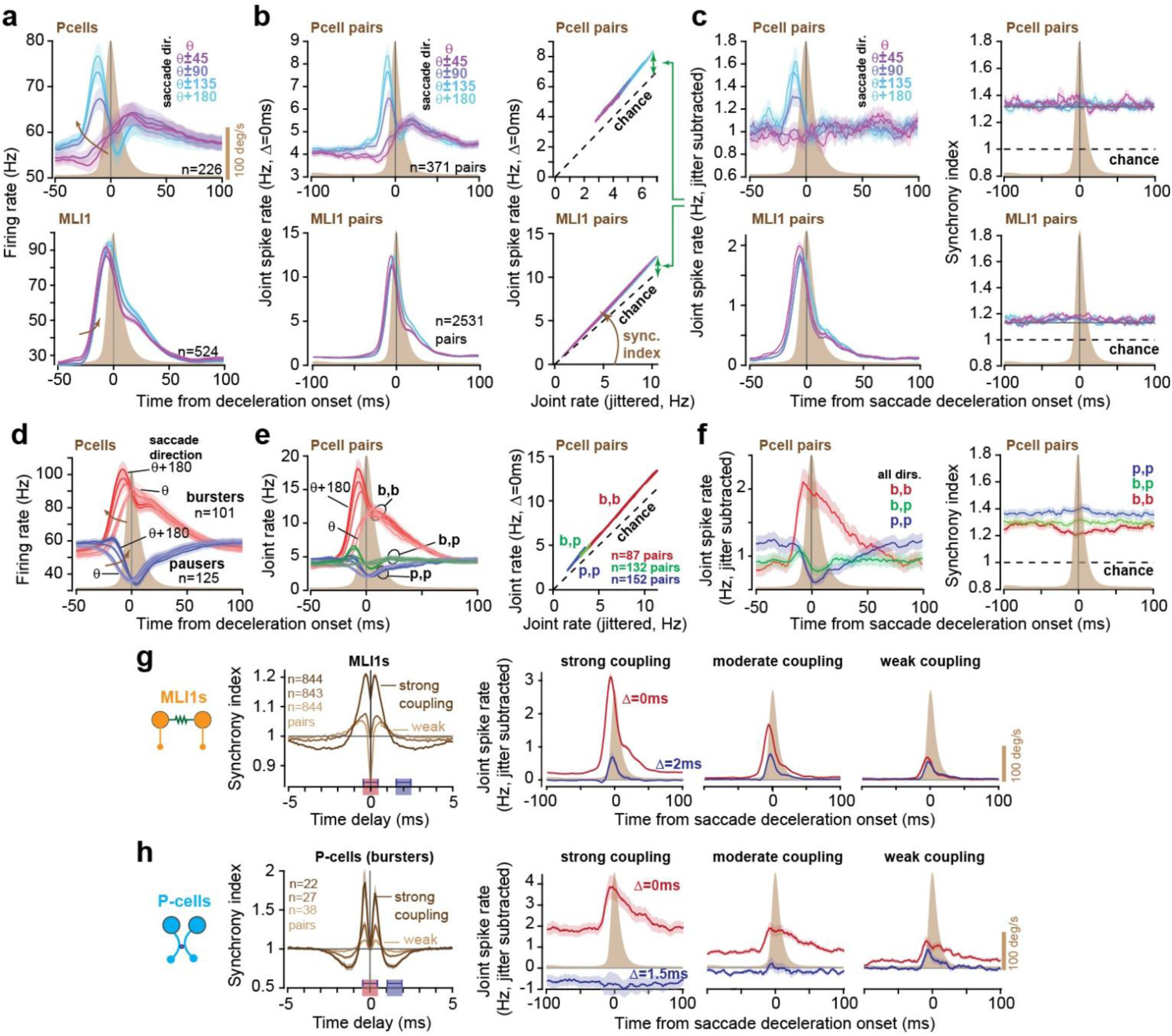
Synchronous spikes were promoted during saccades while asynchronous spikes were suppressed or remained at chance. Data from the saccade periods only. **(a)** P-cell (top) and pMLI1 (bottom) average firing rates aligned to saccade deceleration onset for different movement directions (aligned to each clique’s potent direction). **(b)** (Left) Same as (a) but showing the average joint spike rate at Δ = 0ms delay for each P-cell pair. (Right) Joint-jitter plots for all P-cell (top) and pMLI1 (bottom) pairs from -100 to 100ms relative to saccade deceleration onset, shown separately for each saccade direction (aligned to the clique’s potent direction). **(c)** Average jitter-subtracted (covariance; left) and synchrony index (right) for all P-cell (top) and pMLI1 (bottom) pairs within a clique, aligned to saccade deceleration onset. **(d-f)** Same as in (a-c) but separately for burster and pauser P-cell groups. **(g-h)** Grouping of pMLI1s and P-cells based on strength of electrical coupling. (Left) Synchrony index as a function of delay grouped by strength, 1 is chance. (Right) Joint spike rate at various delays during saccades for groups of strong, medium, and weakly coupled pairs. Error bars are SEM.

Notably, during saccades in direction *θ* + 180, the rate of synchronous spikes above chance was 4.5 times greater if the P-cells were within a clique as compared to between cliques ( peak 9ms before saccade deceleration onset, mean±SEM within: 1.61±0.12Hz vs. between: 0.36±0.04, Fig. 3c & Extended Data Fig. 6G). This is because if the P-cells belonged to different cliques, there was still an increase in the rate of synchronous spikes during saccades, but that increase was almost entirely due to the change in firing rates. Among the pMLI1s, the rate of synchronous spikes above chance was 2.2 times greater within a clique (5ms before saccade deceleration onset, mean±SEM within: 1.86±0.07Hz vs. between: 0.83±0.03Hz).

During a saccade some P-cells increased their rates while others decreased their rates, i.e., bursters and pausers (Fig. 3d). Thus, our cell pairs within a clique consisted of three groups: burster-burster (bb, n=87 pairs), burster-pauser (bp, n=132 pairs), and pauser-pauser (pp, n=152 pairs). These numbers implied that the two types of P-cells often coexisted in close spatial proximity, with a bias toward pauser-pauser pairs. The bb group showed an increase in the joint spike rate during saccades, whereas the pp group exhibited a decrease (Fig. 3e, left subplot). However, regardless of whether the P-cells were bursters or pausers, the joint spike rate remained on the same line (Fig. 3e, right subplot) and the slope, i.e., synchrony index, remained constant (Fig. 3f, right subplot).

Because saccades are only 30-40ms in duration, we were concerned that our ability to measure synchronous rates may be limited. To verify the robustness of our findings, we confirmed that for any pair of neurons, their synchrony index as computed during saccades was the same as the synchrony index computed during the entire recording (Extended Data Fig. 6).

We next considered asynchronous spikes, i.e., 1.5-2ms apart. We stratified the neurons based on their strength of coupling (Fig. 3g & 3h) and found that in pMLI1 pairs with strong coupling, a rise in synchronous rates accompanied only small changes in asynchronous rates, but in pairs with weak coupling the synchronous and asynchronous rates matched each other. In P-cells with strong coupling, asynchronous rates were suppressed below chance during saccades, whereas in P-cells with weak coupling the two rates nearly matched.

Next, we asked whether variations in synchrony produced changes in the animal’s behavior. We identified saccades during which there were more synchronous spikes among a pair of P-cells within clique than predicted from their synchrony index (Extended Data Fig. 7). This division of saccades produced trials in which the average firing rates of the two P-cells in the clique were similar, but the joint rates were different. However, we could not detect any difference in saccade trajectory when the joint rates were larger. That is, presence of greater synchrony (than expected) in a single clique of P-cells did not produce a difference in behavior that we could detect.

In summary, as the firing rates changed during a saccade, the rate of joint spikes (within 0.5ms) peaked near deceleration onset. For P-cell bursters, the joint rate was ∼15Hz. In P-cell and pMLI1 pairs, an increase in firing rates not only increased the rate of synchronous spikes but also suppressed or kept at chance the rate of asynchronous spikes. These patterns demonstrated that spike coordination was not different during saccade and non-saccade periods, hinting that coordination was due to biophysical and anatomical properties of the neurons, something that we will explore shortly.

### Synchronous spikes produced superposition, but asynchronous spikes caused interference

Did production of synchronous spikes in the upstream neurons make a difference in the downstream target? Was production of asynchronous spikes counterproductive? To answer these questions, we began by considering the effects of a single pMLI1 spike upon a P-cell (n=2025 pairs within clique, n=3342 pairs between cliques) by computing the probability of a P-cell spike at time *t* + Δ, given that the pMLI1 produced a spike at time *t* (Fig. 4a, left). We then jitter corrected this probability and found that the inhibitory effect of the pMLI1s was bimodal: after the pMLI1 spike there was an initial period of P-cell inhibition that peaked at a latency of 0.4ms, and then a second period of inhibition that peaked at a latency of 1.3ms. The timing of the initial peak was consistent with electrical interaction at the pinceau, and the timing of the second peak was consistent with chemical interaction via GABA ^21^.

**Figure 4.**
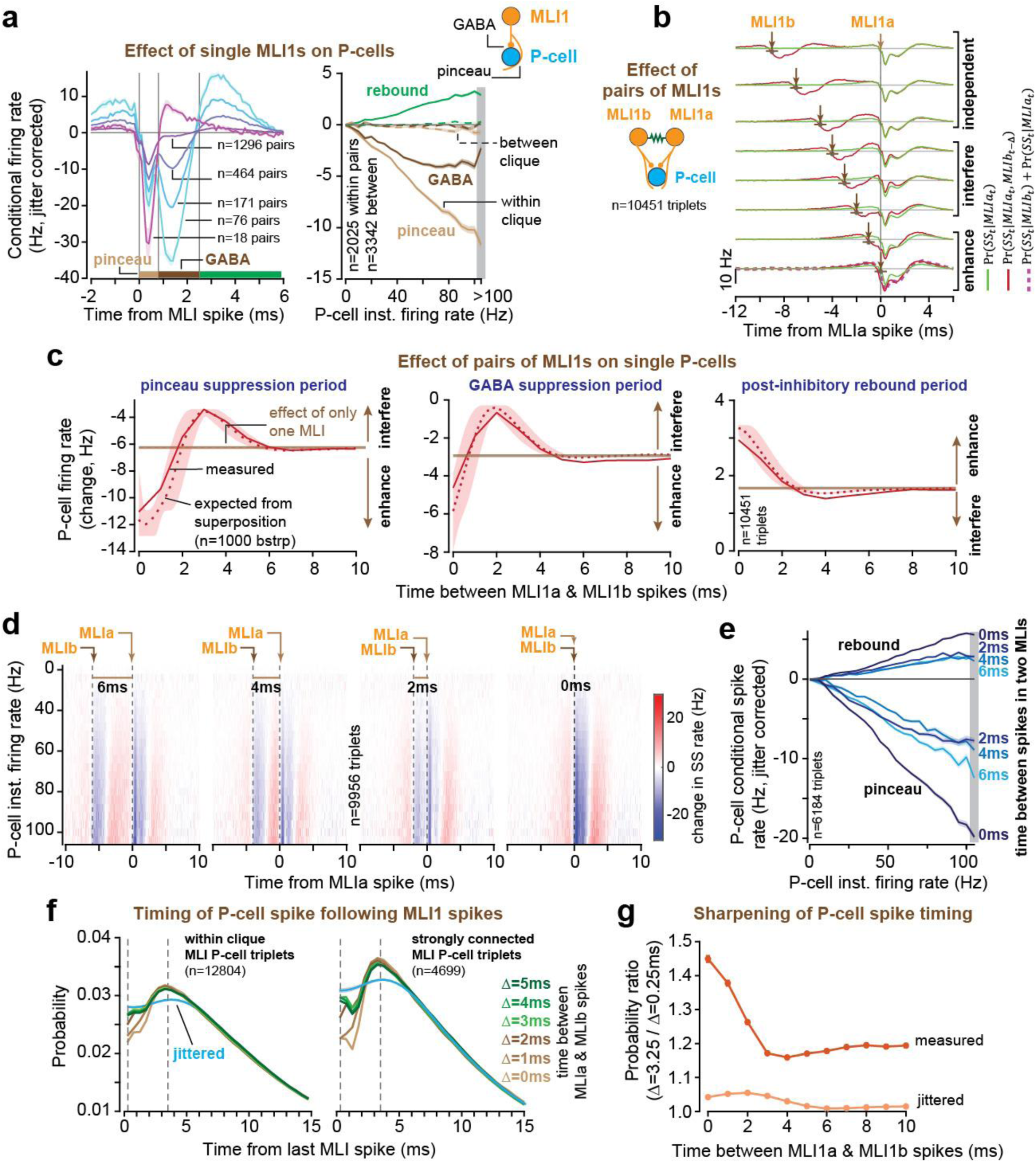
Synchronous pMLI1 spikes induced superposition on the target P-cell, improving precision of simple spike timing. **(a)** (left) Conditional firing rate of P-cell simple spike given a pMLI1 spike (within clique), revealing varying strengths of pinceau, GABA, and rebound. (Right) Average jitter-corrected conditional firing rate for each phase (light brown: pinceau, dark brown: GABA, green: rebound) as a function of the P-cell’s instantaneous firing rate, grouped by within-clique (solid) and between-clique (dashed) pairs. **(b)** Average effect of two pMLI1 spikes on a P-cell at different inter-spike intervals. The green trace shows the effect of pMLI1a alone, red trace is the effect of both pMLI1 spikes, and the dashed red trace shows the linear sum of individual effects from pMLI1a and pMLI1b. **(c)** Triplet analysis. Effects of spikes in pairs of pMLI1s on their target P-cell as a function of their temporal distance. Dotted line is the expected value if the effects of the two spikes summed linearly. The y-axis is the change in firing rate in the P-cell. The horizontal line is the expected effect on the P-cell rate if only one MLI had spiked. The solid red line is the measured effect, i.e., Pr(*SS_T_* |*a_t_*, *b_t_*_+Δ_). The dotted red line is the expected effect due to superposition, i.e., Pr(*SS_T_*|*a_t_*) + Pr(*SS_T_* |*b_t_*_+Δ_). (left) The pinceau period, *T* = (*t* + 0) *to* (*t* + 0.8) ms period. (middle) The GABA period, *T* = (*t* + 0.8) *to* (*t* + 2.5) ms period. (right) The post-inhibitory rebound period, *T* = (*t* + 2.5) *to* (*t* + 6) ms period. **(d)** Triplet analysis. The effects are binned by the P-cell’s instantaneous firing rate. **(e)** Triplet analysis. Pinceau suppression and rebound average amplitudes as a function of P-cell instantaneous firing rate for different delays between spikes in pairs of pMLI1s. Rebound is largest when the two MLI1 spikes are synchronous. **(f)** Triplet analysis. Probability density of a simple spike, following spikes from two pMLI1s. Left: all triplets within a clique. Right: for the triplets with the strongest MLI1 inhibition of the P-cell. **(g)** Sharpening of the probability density of simple spikes, measured as the ratio of the probability of simple spike at 3.25ms to 0.25ms (dashed lines in part f). Error bars are SEM, except for 4c, where the shaded area is the 95% confidence interval for expected traces, obtained by bootstrapping (n=1000).

Some pMLI1s had a strong pinceau effect on a P-cell, some had a strong GABA, and some had both ^34^. There were no significant correlations between the strength of pinceau and GABA within a clique (corr=0.03, p>0.2, Extended Data Fig. 8). Notably, the P-cell inhibition was followed by a rebound starting at 2.5ms (Fig. 4a, left, green horizontal bar). The strength of pMLI1 interactions with P-cells was indistinguishable among P-cell bursters and pausers (Extended Data Fig. 8, pinceau period, burster P-cells: -6.17±0.21Hz vs. pauser P-cells: - 6.44±0.19Hz, p=0.34, GABA period, burster: -3.2±0.18Hz vs. pauser: -3.4±0.17Hz, p=0.43, rebound period, burster: 1.8±0.07 vs. pauser: 1.79±0.06, p=0.89).

These results relied on conditional probabilities, which did not measure the inhibitory current induced by an MLI1 spike on a P-cell. For example, when a P-cell was firing slowly, the effect of a single pMLI1 spike would appear smaller than when the P-cell was firing rapidly. To account for the state-dependent effect of pMLI1s on the P-cells, we adapted the 3D auto-correlogram approach ^35^ and calculated the conditional probabilities as a function of the P-cell’s instantaneous firing-rate (Fig. 4a, right). This revealed that when the P-cells were firing strongly, the effect of the pMLI1 spike was greater, producing a large initial inhibition, followed by a larger subsequent rebound.

We next considered triplets of neurons (n=10,451 triplets) composed of two pMLI1s, termed MLI1a and MLI1b, and one P-cell (Fig. 4b). We organized this data based on time Δ between two consecutive pMLI1 spikes: a spike in MLI1b at time *t* − Δ followed by a spike in MLI1a at time *t*. We then measured the effects that these events had on the simple spikes. For example, consider when the two spikes were more than 6ms apart (Fig. 4b, top row). The green line shows the result when only MLI1a spiked, and the red line shows the result when both MLIs spiked. Each spike inhibited the P-cell, but their effects did not add because the events were too far apart. As the two pMLI1 spikes occurred closer together, their combined effects began to sum so that when they spiked synchronously, the joint event roughly doubled the effect of a single event (Fig. 4c, compare horizontal brown line with red line at 0ms). However, when the two spikes were 2-4 ms apart, the result was interference because the inhibition from the latter spike overlapped with the rebound from the former. Thus, synchronous spikes produced enhancement whereas asynchronous spikes produced interference.

Because the effect of the pMLI1 spikes depended on the state of the P-cell, we computed the state of the P-cell via its instantaneous firing rate and found that for higher P-cell firing rates, the synchronous spikes in the pMLI1s exhibited a larger inhibition, followed by a larger rebound (Figs. 4d & 4e). When P-cell firing rates were around 100 Hz, a rate attained by the bursters during saccades, a synchronous spike in a pair of MLI 1s doubled the probability of P-cell suppression (Fig. 4e, slope of -0.19± 0.002 vs. −0.09 ± 0.002; p=8.7×10^−67^, linear mixed effects), and also doubled the probability of a rebound simple spike, as compared to when the two MLI1 spikes were 4ms apart (slope of 0.06 ± 0.001 vs. 0.03 ± 0.001; p=3.5×10^−54^, linear mixed effects).

In summary, when the spikes in a pair of inhibitory neurons were synchronous, their effect on the downstream target was superposition of inhibition followed by superposition of rebound, but when the two spikes were asynchronous, the result was interference in both the inhibition and the subsequent rebound.

### Synchronous pMLI1 spikes reduced the variance of spike timing in the P-cells

Synchronous pMLI1 spikes were more effective in producing a rebound spike in the downstream P-cell than asynchronous spikes (Fig. 4e). This hinted that if two MLI1s generated a synchronous spike, they could better control spike timing in the downstream P-cell.

To test this idea, we considered triplets in which two pMLI1s converged onto a single P-cell and measured the probability density of a P-cell’s next spike following a spike in each of the two pMLI1s (Fig. 4f). This density exhibited a peak at 3.5ms following the synchronous pMLI1 spikes (Fig. 4f, Δ = 0ms). We then varied the time between the two pMLI1 spikes and found that as Δ became larger, the region around the peak density became broader. To quantify the sharpness of the peak of the density, we took the ratio of the probability of simple spike at Δ = 3.25ms to Δ = 0.25ms and found that this ratio peaked when the time between the spikes generated by the two pMLI1s was 0ms (Fig. 4g). Thus, synchronous spikes in the upstream pMLI1s reduced the variance of the next spike in the downstream P-cell.

### Spike timing exhibited regularity in P-cells but not MLI1s

In individual P-cells, spike timing tended to be “regular” ^11–14^, meaning that each P-cell roughly relied on an internal clock to generate its spikes. To illustrate this, we plotted the 3D-autocorrelogram ^35^, i.e., the conditional probability of spiking at time *t* + Δ, given that the cell spiked at time *t*, organized by the instantaneous firing rate of that neuron (Fig. 5a). The curtain-like bands indicate that each P-cell tended to spike after a duration inversely proportional to its instantaneous firing rate. In contrast, this pattern was not present in pMLI1s. In the P-cells, the probability peaks became sharper as firing rates increased, indicating greater regularity (Extended Data Fig. 9B) ^36^. Did regularity enhance coordination of spike timing?

**Figure 5.**
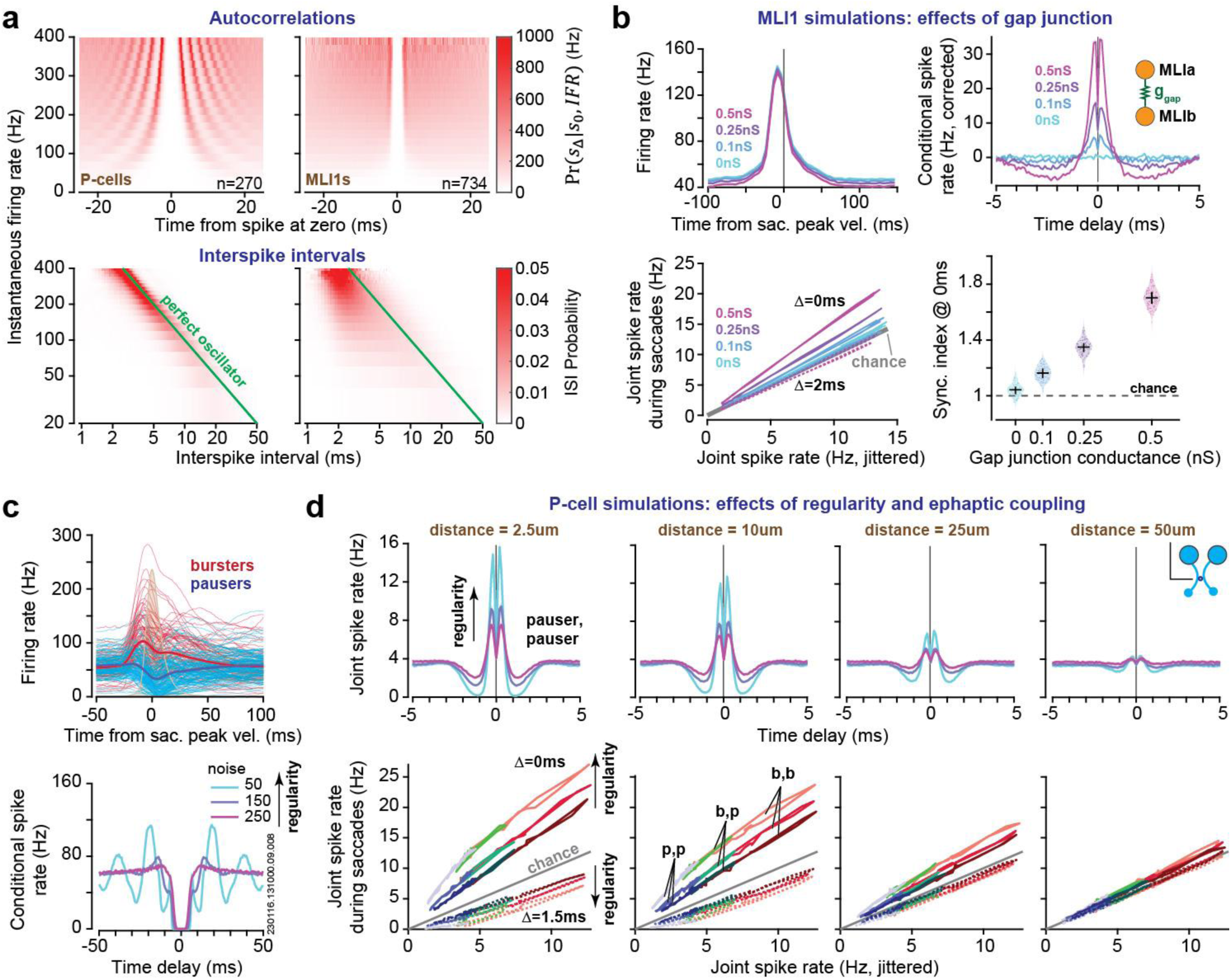
Regularity of P-cells enhanced the production of synchronous spikes. **(a)** Auto-correlograms (top) and ISI distribution (bottom) of all P-cells (left) and pMLI1s (right), plotted as a function of instantaneous firing rates. **(b)** Simulations of pairs of MLI1s during saccades. We varied the strength of gap junction coupling. Top left: average firing rate s during saccades. Top right: conditional spike rate, jitter corrected. Bottom left: joint-jitter plots. Bottom right: Synchrony index (slope of the joint jitter plot) increases with gap junction conductance. Dots indicate individual simulations (n=120). Error bars: median ± interquartile range. **(c)** Top: trial-averaged actual simple spike firing rates for burster P-cells (n = 101) and pauser P-cells (n=125) aligned to saccade peak velocity. The thin lines show the rates of individual P -cells, and the thick line shows the population-averaged saccade response. The average saccade velocity profile is represented by the brown patch. Bottom: auto-correlograms of an example simulated P-cell (a burster), simulated with various regularities. The ID number refers to the actual P-cell firing rates from which the simulated results were produced. **(d)** Simulations of pairs of P-cells during saccades. Top row: joint firing rates for simulated P-cell pairs (bursters) for various strengths of ephaptic coupling and regularities. Bottom row: joint-jitter plots of simulated P-cell pairs (pair type denoted by color: red for burster-burster pairs, green for burster-pauser pairs, blue for pauser-pauser pairs) with different strengths of ephaptic coupling and regularities during saccade (from -50 to 100ms relative to saccade peak velocity). Error bars are SEM.

### Regularity increased synchrony among simulated P-cells

We used simulations to test whether electrical coupling was sufficient to generate the linear joint-jitter relationships in our data, and then determined the roles that gap junctions, ephaptic coupling, and regularity played in coordinating spike timing.

We modeled pairs of MLI1 neurons that had conductance-based gap-junctions and received parallel fiber inputs via granule cells, which in turn received inputs from 3-5 mossy fibers (Fig. 5b). The data for the mossy fibers were from 222 neurons (“state” mossy fibers) recorded during saccades ^10^. As gap-junction strength increased, so did the double-peak structure of the cross-correlograms (Fig. 5b, top row, right column). Without gap-junctions, the synchronous spike rates were at chance (Fig. 5b, bottom row, left column). As MLI1 firing rates increased, the rate of synchronous spikes ( Δ = 0ms) rose above chance while the rate of asynchronous spikes (Δ = 2ms) stayed near chance. Thus, gap-junctions between two simulated MLI1s reproduced the pattern of spike timing that we had recorded in pairs of pMLI1s.

Unlike pMLI1s, which increased their rates during saccades, P-cells displayed heterogeneous activity (Fig. 5c). We simulated P-cells using a leaky integrate-and-fire model that controlled the average firing rate of each cell so that it reproduced the recorded data in terms of bursting or pausing during saccades. We included the ability to set the regularity in the spike timing of individual P-cells via a noise term (Fig. 5c), as well as the strength of ephaptic coupling between pairs of P-cells (Fig. 5d).

We simulated two P-cells with combinations of bursting or pausing activity, and different spike regularities, generating spike trains and quantified their temporal coordination using cross-correlograms and joint-jitter plots (Fig. 5d). Focusing on pairs of bursters, we found that ephaptic coupling enhanced spike synchrony at Δ = 0ms while suppressing asynchrony at Δ = 1.5ms (Fig. 5d, top row, effect of distance). Notably, increased regularity magnified these two features (Fig. 5d, top row, effect of regularity). Fig. 5d also illustrates that in the absence of ephaptic coupling, regardless of spike regularity, the joint probability remains at chance, thus explaining why P-cells in disparate cliques did not exhibit synchrony.

Why should regularity magnify the effects of ephaptic coupling? When one neuron spikes, the extracellular voltage changes by a small amount, typically by 0.5 mV at distances of just a few micrometers. Thus, a spike in one P-cell (PCa) can induce a spike in another P-cell (PCb) if the membrane voltage of PCb is within 0.5 mV of the spiking threshold. We will refer to this voltage region as the ephaptic spiking band. The probability that any randomly selected spike in PCa induces a spike in PCb depends on how much time the membrane voltage of PCb spends in the ephaptic spiking band. If PCb is regular, the amount of time spent in the ephaptic spiking band will be roughly proportional to the width of the band (Extended Data Fig. 9G, left). As noise increases, PCb becomes less regular and the proportion of time spent in the ephaptic spiking band plummets (Extended Data Fig. 9G, right).

Thus, the simulations revealed that the gap-junctions between MLI1s were sufficient to reproduce the temporal coordination observed in our recorded pMLI1s, and ephaptic coupling was sufficient to reproduce the coordination observed in our recorded P-cells. The clock-like regularity of P-cells increased the rate of synchronous spikes while suppressing asynchronous spikes (in simulations).

### Ephaptic coupling reset the phase of neighboring P-cells

If P-cells rely on an internal clock, then the time between two consecutive spikes can be viewed in terms of the phase of the clock. We obtained an estimate of the phase by multiplying the time elapsed from each spike by the instantaneous firing rate at the time of that spike. This new way of representing time produced a novel measure of auto correlations in terms of the phase of the clock (Fig. 6a). For the P-cells, but not the pMLI1s, the result was a wave function with peaks at multiples of 2*π*. Thus, the P-cells resembled tunable clocks that could run fast or slow (Fig. 6b).

**Figure 6.**
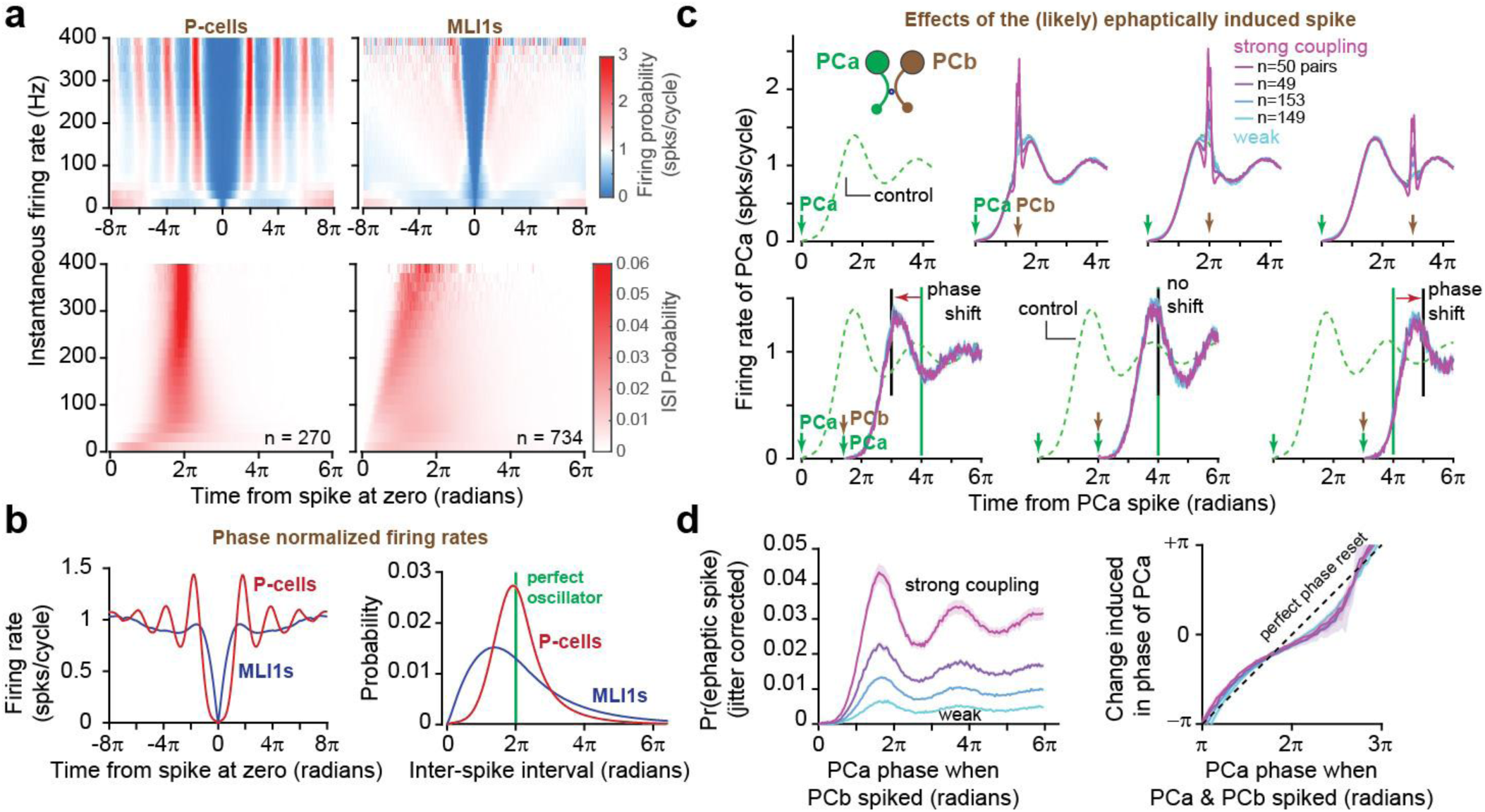
Spike timing of P-cells relies on an internal clock. Ephaptic coupling allows one P-cell to reset the internal clock of another P-cell. **(a)** Phase normalized auto-correlograms (top) and ISI distribution (bottom) of all P-cells (left) and pMLI1s (right), plotted as a function of instantaneous firing rates. (b) Mean auto-correlogram (left) and ISI distribution (right) of P-cells and pMLI1s in phase domain. (c) Top: mean conditional probability of PCa firing, given that a neighboring PCb spiked at a phase relative to PCa: from left to right: control (jittered PCb spike), PCb at 7*π*/5, 2*π*, and 3*π*. Bottom: mean probability of PCa firing given that PCa fired again less than 0.5 ms after a PCb spike occurring at some phase Δ relative to PCa (from left to right Δ = 7*π*/5, 2*π*, and 3*π*) grouped by different PCa and PCb ephaptic coupling strength. Note that the spike in PCb resets the phase of PCa. **(d)** Left: mean probability of ephaptic spike (jitter corrected joint spiking of PCa and PCb) given that PCb spiked at different phases of PCa, grouped by different PCa and PCb ephaptic coupling strength. Right: circular mean of the change in phase of PCa as a function of when the ephaptic spike occurred. Shaded areas are SEM for all figures, except for 6D right, where the shaded area is the 95% confidence interval on the circular mean, obtained by bootstrapping on the population of P-cell pairs.

Now imagine that the clock of a P-cell (PCa) is running and has reached *π*, i.e., half of its cycle. At this time PCb interrupts PCa’s clock by ephaptically generating a spike, moving PCa’s phase to 2*π*. Does this interruption reset the clock of PCa? For each P-cell within a clique we plotted its normalized firing rate as a function of phase (Fig. 6c, green dashed line). When PCb generated a spike, it increased the probability of spiking in PCa (Fig. 6c, top row). Occasionally, PCa spiked at less than 0.5ms after PCb. To test whether a joint spike could reset PCa’s phase, we grouped P-cell pairs by the strength of their ephaptic coupling (synchrony index, Fig. 2f) and examined whether the joint spike acted like a regular spike in resetting the clock. When the joint spike occurred before 2π, the subsequent spiking probability retained the same oscillatory nature, but with a phase shift relative to control, so that the peaks in spiking probability now occurred earlier. This effect was consistent across all levels of ephaptic coupling strength, suggesting that the joint spike reset the phase no differently than spontaneous ones (Fig. 6c, bottom row). When the joint spike occurred after 2π, a phase delay was induced in the spiking probability, so that the peaks were now later than the peaks in the control. In contrast, if the joint spike occurred near 2π, there was minimal shift in the conditional spiking probability.

We next estimated the probability of an ephaptically induced spike through subtraction of joint spike rates from chance (jittered PCb spikes). P-cells with stronger couplings showed higher probabilities of estimated ephaptic spikes for all phases of PCa phase (Fig. 6d, left). Spikes from a neighboring P-cell were most effective at inducing ephaptic spikes in PCa when the phase of PCa was close to a multiple of 2π (i.e. when PCa was already close to firing), regardless of ephaptic coupling strength (Fig. 6d, left). Additionally, regardless of the strength of their coupling, the change induced in the phase of PCa by the joint spike was roughly a linear function of when that spike occurred (Fig. 6d, right). For example, if this spike occurred before PCa would have normally spiked (before 2π), it moved the next PCa spike to an earlier phase in the cycle, and if it occurred after PCa would have normally spiked (after 2π), it moved the next PCa spike to a later phase in the cycle. To check whether these results were trivial, we confirmed that pMLIs did not show evidence of phase resetting (Extended Data Fig. 9).

Thus, when one P-cell induced a spike in its neighbor, the synchronous production of spikes aligned the phases of the two clocks. This phase alignment produced a phase reset in the target P-cell and shifted its next spike timing. If the instantaneous firing rate of the two P-cells were similar, the phase alignment might increase the likelihood of next spike being synchronous as well.

## Discussion

We discovered that spike timing in two types of electrically coupled inhibitory neurons, P-cells and pMLI1s, obeyed a mathematical pattern: as the firing rates of individual neurons increased, the rate of joint spikes that were separated by a given time interval rose linearly such that synchronous intervals were promoted while asynchrony was suppressed. To understand the consequences of this coordination, we isolated pairs of pMLI1s that converged on a target P-cell. When the upstream pair spiked synchronously, the downstream neuron experienced a doubling of the inhibition, then a doubling of the post-inhibitory rebound. However, when the upstream spikes were 2-3ms apart, the result was mutual interference.

Thus, electrical coupling promoted spike intervals that produced constructive superposition upon the target while suppressing intervals that produced destructive competition. The result was better control of spike timing in the target neuron: synchronous spikes reduced the variance of the next spike in the downstream neuron.

Spike timing of P-cells exhibited a clock-like pattern such that each P-cell resembled a tunable oscillator. When a P-cell ephaptically induced a spike in its neighbor, that event aligned the phases of the two oscillators. Simulations suggested that this clock-like conformity may increase the efficacy of P-cell ephaptic coupling, resulting in an increased number of joint spikes.

### Proximity and biophysics largely accounted for coordination of spike timing

The spike timing of pMLI1 pairs were consistent with gap-junctions ^3,37^, producing increased synchronous rates as firing rates increased ^38^. Our simulations of gap-junctions in the MLI1s and ephaptic coupling in the P-cells reproduced the linear relationship between joint and jittered rates (Eq. 1). This suggested that electrical coupling and regularity were sufficient to account for spike timing coordination.

But does this mean that the cerebellum does not have the ability to change synchronous rates beyond those dictated by its biophysics and anatomy? Dopamine can rapidly alter the strength of gap-junctions ^39^, and both P-cells and MLIs have dopamine receptors ^40^. Furthermore, the regularity of a neuron’s spike timing is not fixed and can be altered in short timescales ^41,42^. Thus, neuromodulatory and other inputs to the cerebellum may have the means to modify coordination of spike timing.

### Joint-jitter plots resolved controversies in measuring coordination of spike timing

How should one examine whether the rate of synchronous events is different than chance? One method measures the ratio of joint probability to the independent probabilities, termed the synchrony index ^43^. However, as the firing rates decline to near zero, this ratio becomes noisy or undefined, raising concerns regarding our observations in a previous work ^9^. To remedy this, Herzfeld et al. ^8^ suggested measuring the difference between the rate of synchronous events and the independent probabilities, termed covariance. However, the joint rate can be an arbitrary function of the firing rates of the two neurons, making it unclear whether synchrony index or covariance is the proper measure.

We solved this problem by quantifying the empirical relationship between joint and independent probabilities. The resulting joint-jitter plots revealed that the joint rates varied linearly as a function of the jittered rates, meaning that synchrony index remained constant.

### Ephaptic coupling and clock-like regularity combined to increase synchrony

What was the functional significance of clock-like regularity in P-cell spike timing? Payne et al. ^36^ investigated this question by controlling the timing of spike production in groups of P-cells via optogenetic stimulation. They demonstrated that each pulse of stimulation moved the eyes by a small amount, but this effect was not dependent on whether the stimulation timing was regular or irregular. Here, our simulations revealed that regularity works in concert with ephaptic coupling, predicting that P-cells that exhibit greater regularity ^44–46^ will also exhibit stronger synchrony. Thus, a functional purpose of regularity may be to promote near synchronous spike time intervals while suppressing asynchronous intervals.

However, we could not detect changes in saccade kinematics when a pair of P-cells generated more synchronous spikes than expected from their joint-jitter relationship (Extended Data Fig. 7). Moreover, we do not know the extent to which P-cells remain bound to their internal clocks during behavior. This is because it is difficult to estimate instantaneous firing rates during single saccades.

### A rationale for coordination of spike timing among P-cells

Whereas synchronous pMLI1 spikes produced superposition on the target P-cell, spikes that were but a few milliseconds apart were destructive because, rather than adding to one another’s contributions, they reduced these effects. Roughly 50 MLI1s converge onto the pinceau of a single P-cell ^47^, and around 50 P-cells converge onto a single nucleus neuron ^15^. We found that like pMLI1s, P-cell pairs strongly promoted joint spiking within 1ms of each other but suppressed intervals of 2ms. The inhibitory post-synaptic current produced by a single P-cell spike in the nucleus neuron has a time constant of 2.5ms ^15^. This extremely fast time constant implies that, like the effect of the MLIs on the P-cells, only synchronous P-cell spikes can induce superposition in the nucleus neuron while 2ms spike intervals cause destructive interference.

Gauck and Jaeger ^48^ found that spiking in the nucleus neurons can be driven by the production of synchronized P-cell inputs, or a transient reduction in their firing rates. However, an increase in the inhibitory input had little effect. Nashef et al. ^49^ confirmed this using a dynamic clamp experiment in nucleus neurons, demonstrating that nucleus neurons increased their spiking under two scenarios: when P-cell firing rates declined, and synchrony increased. This implies that the cerebellar cortex combines two mechanisms to drive the activity of the nucleus neurons: reduced inhibition via pausing P-cells, and increased synchrony via bursting P-cells.

Whereas synchronous spiking among P-cells has been shown to entrain downstream nucleus neurons ^15,48,50^, a recent work found that synchronous pausing, i.e., not spiking for relatively long periods, also has a causal effect on nucleus firing ^51^. Indeed, there is evidence that synchronous pausing is also regulated among pairs of P-cells ^52–55^. In the zebrafish cerebellar nucleus, the activities of the nucleus-like neurons cannot be reproduced via mean rates of inhibitory and excitatory inputs, because on average, mean inhibition far exceeds excitation. However, firing rates in the nucleus depend on precise timing of the inhibitory and excitatory inputs, allowing for gaps in inhibition to bring the neuron to threshold ^51^.

Thus, both the rate and the timing of P-cell spikes, including synchrony and coordinated pauses, appear critical for shaping the output of cerebellar nucleus neurons.

### The role of MLIs in coordinating spike timing in pairs of P-cells

Parallel fibers provide excitatory inputs to both the P-cells and the MLIs, resulting in a feedforward inhibitory network in which inhibition can rapidly quench the excitation ^56^. The MLI1s employ the pinceau to generate one of the fastest inhibitions in the entire brain ^21^. The electrical coupling between MLI1s appears to make them responsive to synchronous events in the parallel fibers ^57^, raising the possibility that the role of the feedforward inhibitory network is to make P-cells respond mainly to coincident events in their parallel fibers. In addition, it is possible that a component of spike coordination among pairs of P-cells may be due to a network wide organization that involves the MLI1s ^58^. However, without the GABA induced inhibition from stellate cells ^11^, or with gene knockout of P-cell GABA receptors ^14^, P-cell spike timing becomes more regular.

### Spatial organization of temporal coordination

Clustering neurons into cliques was required for unmasking the organization of spikes ^10^, but the spatial extent of a clique remains unknown. Here, each clique was confined to a single P-cell layer, and its width may extend along the parallel fibers ^59^. Previous work suggests that P-cell to P-cell interactions are not random but spatially structured ^5,60^. The connectivity matrix of P-cells (Extended Data Fig. 2) varies across cerebellar regions ^61^. Recordings that span larger areas of the cerebellar cortex using multi-shank probes will be essential to reveal the network properties arising from these couplings.

### Cerebellar disease typically accompanies disruption of spike timing, not firing rates

Genetic mutations that disrupt function of calcium channels generally do not affect average firing rates of P-cells, but instead alter their clock-like regularity ^62^. In Duchenne muscular dystrophy, P-cell firing rates remain stable while regularity is altered ^63^. In episodic ataxia type-2, P-cells exhibit reduced regularity ^64^, and the drug 4-AP improves the symptoms while restoring P-cell regularity ^65^. However, regularity may not encode any information beyond what is already present in the firing rates ^36^. Rather, regularity combined with ephaptic coupling may increase synchrony in P-cell pairs, which in turn may enhance the ability to control timing of nucleus neuron spiking.

### Implications beyond the cerebellum

The principles revealed here may apply broadly to cerebral cortex and hippocampus, regions that contain fast-spiking interneurons that are electrically coupled ^1,66^. Our results suggest that electrical coupling will bias spike timing toward intervals that maximize temporal summation in downstream targets while minimizing interference, thereby improving the precision of spiking in the target neurons.

A related principle may apply to the retina where neighboring neurons are electrically coupled ^67,68^. The synchronous spikes that these cells produce improve encoding of visual features and increase reliability of transmission to downstream structures ^69^. Our results suggest that electrical coupling provides control of spike timing in the downstream thalamic neurons.

Overall, we found that electrical coupling provided inhibitory neurons with the means to improve the precision of spike timing in their downstream targets. Clock-like regularity, a feature that is present in P-cells and other electrically coupled inhibitory cells such as thalamic reticular neurons ^2^, is likely to strengthen this control.

## Acknowledgements

We are grateful to Paul Hage who helped with surgeries and recordings, and Jay Pi who helped with writing the behavior software and setting up the recording chamber. We are grateful to all members of Shadmehr Lab for their input and suggestions with this project and help with care of the animals. We are grateful to Prof. Tim Harris for the Neuropixels Ultra probes.

## Author contributions statement

MAF and AMS recorded all electrophysiological data. MAF, EAT, and ANV analyzed the data and made figures. ANV and MAF performed neural modeling. EAT and MAF devised algorithms for measurement of synchrony. EAT devised algorithms for measurement of regularity and phase resetting. RS wrote the paper with assistance from all the authors.

