## supplementary file for "Coordination of spike timing among the neurons of the cerebellum"

### Methods

We collected data from two marmosets, *Callithrix Jacchus*, 350-370 g, subjects 132F (Charlie, 7 years old), and 65F (Barney, 7 years old), during a 2-year period. The marmosets were born and raised in a colony that Prof. Xiaoqin Wang has maintained at the Johns Hopkins School of Medicine since 1996. Our procedures were approved by the Johns Hopkins University Animal Care and Use Committee in compliance with the guidelines of the United States National Institutes of Health.

*Data acquisition*

Following recovery from head-post implantation surgery, the animals were trained to make saccades to visual targets and rewarded with a mixture of applesauce and lab diet [70]. Visual targets were presented on an LCD screen with 500Hz refresh rate and low latency (Dell AW2524H). Binocular eye movements were tracked at 2000 Hz using VPIXX eye tracking system. Tongue movements were tracked with a 522 frame/sec Sony IMX287 FLIR camera, with frames captured at 100 Hz.

We performed MRI and CT imaging on each animal and used this data to design an alignment system that defined trajectories from the burr hole to various locations in the cerebellar vermis [70], including points in lobule VI and VII. We used 3D Slicer software to first align the T2 MRI to the marmoset atlas [71]. We then aligned the CT image to the T2 MRI. We used a piezoelectric, high precision microdrive (0.5 micron resolution) with an integrated absolute encoder (M3-LA-3.4-15 Linear smart stage, New Scale Technologies) to advance the electrode. To reach the cerebellar cortex, we planned a posterior burr hole and avoided the confluence of sinuses using MRI T2 images. This approach enabled us to record from multiple folia in lobules VI and VII simultaneously.

We recorded from the cerebellum using Neuropixels 1.0, Neuropixels Ultra, as well as 64-channel Cambridge Neurotech probes (M1 and M2). For Neuropixels we used the National Instrument acquisition system (NI PXIe-1071). Data was sampled at 30 kHz and aligned to the eye tracking system time as the reference time using a random TTL signal. For the Cambridge probes we connected each electrode to a 64-channel head stage amplifier and digitizer (RHD2132 and RHD2164, Intan Technologies, USA), then connected the head stage to a communication system (RHD2000 Evaluation Board, Intan Technologies, USA). Because the conductive coating on the Cambridge probes degraded after each insertion into the brain, we re-coated the probes and restored their low impedance after every 3-4 recording sessions [72]. For accurate reaction time and visual target time measurements, we measured the video monitor's latency in real-time with a photodiode and aligned it to the reference time.

*Behavioral protocol*

Each trial began with fixation of a center target after which a primary target appeared at one of 8 randomly selected directions at a distance of 5-6.5 deg. As the subject made a saccade to this primary target, that target was erased, and a secondary target was presented at a distance of 2.5-3.5 deg, also at one of 8 randomly selected directions (Fig. 1a). The subject was rewarded if following the primary saccade, it made a corrective saccade to the secondary target, landed within a 1.5-2.0 deg radius of the target center, and maintained fixation for at least 200ms. The reward was food that was provided in two small tubes (4.4 mm diameter), one to the left and the other to the right of the animal. A successful trial produced a food increment in one of the tubes and would continue to do so for 50 consecutive trials, then switch to the other tube. Because the food increment was small, the subjects naturally chose to work for a few consecutive trials, tracking the visual targets and allowing the food to accumulate, then stopped tracking and harvested the food via a licking bout [29,30].

We analyzed eye movements using a deep neural network that detected all saccades and microsaccades [73]. The pre-trained networks for human and macaque monkeys did not perform well for marmosets. Hence, we designed a Matlab GUI-based program to curate the saccades https://github.com/ShadmehrLCMC/SACCURATE. We then retrained the network for each individual animal using seven 30-45 minutes of recording. To prevent any potential false positive saccades, we pruned the saccades by fitting a bivariable Gaussian distribution to two different feature spaces defined as biologically relevant metrics [74] and removed saccades that were outliers (less than 1% chance of belonging to the distribution). The two feature spaces were log-log main-sequence plots (maximum velocity vs. amplitude) and log-log acceleration time to deceleration time ratio vs. amplitude of saccade. We then analyzed and found valid fixations among candidate fixations after or before each detected saccade.

We detected the onset and offset time of each saccade using a trained neural network, as described above. We low-pass filtered the eye position traces with a 3rd order Butterworth filter with 100 Hz cut-off frequency. We calculated the saccade velocity by differentiating the eye position trace and found peak velocity times and values.

*Neurophysiological analysis*

We used OpenEphys [75] for electrophysiology data acquisition, and then used Kilosort 2.0 and Phy 2.0 [76] to manually identify and curate the spikes. Each recording was curated twice by two experienced neurophysiologists. We controlled for contamination in the cross-correlograms in P-cell simple and complex spikes and removed the spikelets and double detection of CSs as SSs by aligning the two waveforms to each other and removing the copies in SS units [77]. We further tracked the sorted units across three to seven 30-45 minutes of recording using a semi-supervised custom written MATLAB GUI software. To do so, we used electrophysiology properties of the cells including waveform and location on the probe, auto-correlogram and raster plots aligned to saccade onsets. We discarded unstable cells with varying baseline firing rates across time (commonly due to the physical pressure of the probe). Hence not all cells were present for the whole duration of recording. Extended Data Fig. 1C shows the duration of recording for 50 sessions and the distribution of duration of each cell type. For pair analysis we used the overlapping recordings in which both cells were present.

*Quantifying the isolation quality of each neuron*

A summary of the measures used to verify the quality of the neurophysiological data is provided in Extended Data Fig. 1A. To measure the isolation quality of each neuron, we computed the conditional probability $\Pr(s(t+\Delta)=1|s(t)=1)$, that is, the probability of a spike at time delay $\Delta$, for a bin width $w=0.1$ms, given that the neuron produced a spike at time zero. We then multiplied this probability by 10000 (to produce units of spikes per second, Hz) and plotted the results as firing rate (for complex spike auto correlograms, bin width $w = 2$ms). This data is shown separately for confirmed P-cells as well as putative P-cells in Extended Data Fig. 1E.

For well-isolated cells, we expected a clean refractory period. To measure the noise rate in each neuron's spiking, we quantified the average conditional probability at $\Delta \leq 1$ms (except for complex spikes, for which we used a 10ms period). For example, to quantify the quality of the P-cells, we measured the noise rate and found that for the simple spikes, this rate was 0.35±0.26 Hz (median ± median absolute deviation, MAD). For the complex spikes, the noise rate was 0±0 Hz.

We also quantified the violations of simple spike occurrence after complex spikes using their average cross probability for a 5ms period. The data for all cell types are shown in Extended Data Fig. 1A, second row.

*Cell type identification*

We performed manual cell type identification through identification of the layers. First, we identified the P-cells via suppression of SS following a CS. As P-cells are large cells, their spike waveforms are present on multiple channels of the silicon probes (Fig. 1c, Extended Data Figs. 2D & 4). Next, using the CS waveforms on the channels we identified the orientation of the P-cell's axon and dendrites, thus identifying the molecular, Purkinje, and granular layers [16,17,77]. For example, we identified the molecular layer via the downward, broad CS waveforms in the dendrite tree of the P-cells (Fig. 1c). In the granular layer we looked for spike waveforms that exhibited an "m" shape with a slow negative after-wave and labeled those cells as mossy fiber glomeruli [78–80]. In the molecular layer, we labeled neurons that inhibited the P-cells at 1ms latency or sooner as pMLI1. Neurons in the molecular layer that inhibited pMLI1s, excited a P-cell, and experienced excitation from CS spillover were labeled as pMLI2s [18–20,23].

We confirmed that the spatial properties of the spike waveforms (which led to identification of the various layers) were present in both Neuropixels and Cambridge recordings (Extended Data Fig. 3). We also confirmed that the cell-cell interactions were present in both Neuropixels and Cambridge recordings (Extended Data Fig. 3).

Following the identification of the molecular, P-cell, and granule layers, we identified the putative MLI1 and MLI2s (labeled as pMLI1 and pMLI2) based on their spike interactions with each other and the P-cells [20]. These interactions were extracted and measured through cross-correlograms that were then corrected via spike jittering: for each cell pair, the interaction was measured via a cross-correlation, then corrected for chance interaction due to firing rate fluctuations using interval jittering [27], as shown in Fig. 1e, right column. pMLI1s were identified using their milliseconds inhibition of P-cells as well as their synchrony with each other, while pMLI2s were identified through their millisecond suppression of pMLI1s and lack of interaction or later disinhibition of P-cells [20] as shown in Extended Data Fig. 3.

We relied on the interactions of MLIs with each other and with P-cells to identify them, then confirmed that their waveform, baseline rates, and auto-correlograms were similar to previous findings in definitively identified neurons in mice [20]. However, a recently discovered interneuron termed Ito cells (IC) also resides in the molecular layer and inhibits P-cells [81]. Our pMLI1s are likely to exclude these cells because unlike pMLI1s, ICs do not have gap-junctions and do not exhibit spike synchronization.

We identified the mossy fibers based on their unique "m" shaped spike waveform in the granule layer. We did this by relying on our Neuropixels Ultra recordings to acquire a very high resolution "picture" of the mossy fiber spike waveforms (see Supp Fig. S7A of [10]). As previously reported, mossy fibers exhibit a triphasic waveform with a negative after wave or a fast narrow spike [80,82–86]. We used an approach similar to [87] to extract spatio-temporal waveforms of each cell by concatenating spike waveforms on different channels to form a 2D image, then centered the absolute peak to the center of image in time and space (see Supp Fig. S7C of [10]). To get a consistent result for the checkerboard and linear probes we picked two columns out of four columns of the checkerboard probe with highest waveform values for each cell and linearized it. We then centered the absolute peak of the waveform in time and space. Finally, we used UMAP to cluster the mossy fibers from other granular layer interneurons (Supp Fig. S7D of [10]). UMAP on a linearized feature space of these spatio-temporal waveforms revealed two groups of mossy fibers and a potential Golgi cell group. The GLI group (likely composed of mostly Golgi cells), on the other hand, had a larger signature in space and showed wider spike waveforms (Supp Fig. S7E of [10]).

*Clustering of neurons into cliques*

After acquiring the cross-correlograms for each pair of neurons, we jitter-corrected the result [27]. This clustered the neurons into small networks, called cliques.

For each recording session, we computed all pairwise jitter-corrected cross-correlograms. To quantify short-latency interactions between neurons, we examined the 0–3 ms lag window and extracted the maximum absolute interaction value within this interval for each pair (Extended Data Fig. 2E). These values were then used to construct a directed adjacency matrix, $A_{n \times n}$ where n is the number of recorded neurons in that session, and each neuron was a row and a column in the matrix. Some pairs within a session did not have an overlapping recording period; we replaced these values with zeros for spectral clustering.

This adjacency matrix is not necessarily symmetric. However, by the Bayes rule, the cross-correlogram of neuron x conditioned on neuron y and that of neuron y conditioned on neuron x are time-reversed versions of one another and differ only in scaling. To improve robustness, we used 1 ms binning rather than 0.1 ms binning when constructing these matrices. This reduced noise and, importantly, produced single peaks centered near 0 ms for reciprocal electrical interactions such as ephaptic coupling or gap junctions (Extended Data Fig. 2F top). In contrast, directional interactions, such as MLI1 suppression of P-cells or MLI2 suppression of MLI1s, tended to produce asymmetric patterns, with only one of the two transposed matrix elements showing a strong interaction (if any; see Extended Data Fig. 2F, top panel, where Pr(PC|MLI1) shows suppression only on one side of the correlogram).

Because our clustering analysis was intended to capture the strength of interaction between neuron pairs, rather than the directionality of those interactions, we made the adjacency matrix symmetric by assigning each pair (i,j) the larger absolute value of $A_{ij}$ and $A_{ji}$. This produced a symmetric matrix $A_{n \times n}$ representing pairwise interaction strength (Extended Data Fig. 2E).

Similarly, for the purpose of identifying cliques, we were interested in the magnitude of interactions rather than their sign. We therefore defined the weight matrix $W$ as the absolute value of the symmetric adjacency matrix $A$ such that larger positive values indicated stronger interactions (in Hz).

Next, we used spectral clustering to divide the adjacency matrix into groups. Spectral clustering is a graph clustering method which uses the adjacency matrix of the graph and represents the nodes (each cell) in a 2D or 3D subspace of eigenvectors of the Laplacian matrix (spectral space). In the spectral space the interconnected nodes have shorter distance from each other than the weakly connected nodes [88].

We computed the graph Laplacian matrix $L_{n\times n}$, defined for a weighted graph as:

$$L_{n\times n} = D - W$$

where $D_{n\times n}$ is a diagonal "degree" matrix (summation of weights to each node), defined as:

$$D_{ii} = \sum_{j=1}^{n} W_{ij}$$

Next, we performed eigen value decomposition of the Laplacian matrix, where $U_{n\times n}$ contains the Eigen vectors and $\Lambda_{n\times n}$ contains the eigen values:

$$L = U\Lambda U^T$$

The number of zero eigen values corresponds to the number of connected components in the graph. Therefore, $L$ always has at least one zero eigen value. We used the second and third eigen vectors, i.e., the columns of $U$ corresponding to the second and third smallest eigen values as a 2D spectral embedding of the neurons. The representation is shown for an example session in Extended Data Fig. 2F bottom.

As shown in Extended Data Fig. 2F bottom, the two cliques are clearly separated in spectral space, with each clique containing both MLI1s and Purkinje cells. Reordering the rows and columns of the adjacency matrix according to this clustering reveals a block-like structure consistent with two cliques: interactions are stronger within cliques and weaker between cliques (Extended Data Fig. 2E).

Clustering in spectral space was performed manually in multiple steps to separate the main cliques while excluding outliers. A straightforward way to evaluate clustering quality is to compare interaction strengths within and between the identified cliques (Extended Data Fig. 2G top). To illustrate this directly, we also show all high-resolution pairwise interactions for each group as heatmaps in Extended Data Fig. 4B. However, because these values are expressed in Hz, they do not provide a normalized measure of clustering quality across sessions.

We further verified our cell type identification by ordering neurons in each clique in the adjacency matrix by hierarchical clustering. This algorithm organized the rows of the matrix by putting similar rows next to each other. The result was blocks of sub-matrices with similar interactions within and between cell types.

*Validating clique membership*

We used the concept of silhouette values to test the quality of our clustering. For each neuron $i$, we defined the average within clique interaction strength $a(i)$ as the mean weight between neuron $i$ and all other neurons in its clique $c(i)$. We then defined the between clique interaction strength $b(i)$ as the maximum mean interaction of neuron $i$ with neurons in other cliques.

$$b(i) = \max_{c(i)\neq c(j)} \frac{1}{n_j} \sum_{k} W_{ik}$$

We then defined a silhouette-like membership score for each neuron:

$$s(i) = \frac{a(i) - b(i)}{\max(a(i), b(i))}$$

This score ranges from -1 (poor clique membership) to +1 (strong clique membership). The distribution of $s(i)$ across recorded neurons is shown in Extended Data Fig. 2G bottom, with a median ± MAD of 0.72±0.13, indicating strong separation between the identified cliques.

*Finding the Purkinje layer based on complex spike waveforms within a clique*

To further validate our cell-type classification based on the spatiotemporal features of units within each clique, we analyzed the complex spike (CS) waveforms across channels. For each CS unit, we extracted waveforms from its 32 nearest channels and included all of them in the analysis. Channels with absolute peak amplitudes below 25 μV were classified as noise (Extended Data Fig. 2B). We then combined all CS waveforms and applied UMAP for dimensionality reduction followed by Gaussian mixture model (GMM) clustering to separate dendritic and somatic (axonal) CS waveforms.

For each clique, we represented every channel's CS waveform as a point labeled 1 (axonal, sharp initially negative spike followed by a long positive period) or −1 (dendritic, broad negative spike) [16,17]. Estimating the location of the Purkinje layer was then formulated as finding the decision boundary that best separates these two classes. Because the UMAP clustering (Extended Data Fig. 2B) was not perfectly separable and some samples lay between dendritic and axonal clusters, we used a robust classifier to handle label uncertainty. Specifically, we applied a weighted logistic regression model designed to reduce the influence of noisy labels (Extended Data Fig. 2C). Each sample's weight was proportional to its absolute peak amplitude, normalized by the maximum amplitude within its own group (dendritic or axonal) in each clique; in Extended Data Fig. 2C, dot sizes represent these weights.

The resulting decision boundaries were then used to rotate and translate the simple spike (SS) and pMLI1 waveforms to align them with the estimated Purkinje layer in each clique. We further realigned the spatiotemporal waveforms along the time axis to match their spike times (Figs. 1c & 1d). This procedure successfully reproduced the expected simple spike waveforms previously reported in rodent studies [17]. Interestingly, after alignment, pMLI1 waveforms exhibited an upward spike near the P-cell axon—a surprising feature, since upward deflections typically occur near dendrites during simple spikes. However, this pattern was consistent with pinceau recordings in rodents [89]. A representative multi-contact view of two P-cells, two pMLI1s, and the estimated Purkinje layer within a clique is shown in Extended Data Fig. 2D.

*Comparing results of Neuropixels with Cambridge silicon probes*

Our observation of neuronal cliques relied on the ability to record simultaneously from neurons in multiple P-cell layers. Neuropixels had 384 electrodes that spanned ~4mm, while Cambridge M1 had 64 electrodes that spanned ~0.6mm. To check the robustness of our findings, we compared the spike waveforms and spike interactions for simple spikes, complex spikes, MLI1 spikes, and MLI2 spikes, in each probe. The spike waveforms recorded by the two probes are shown in Extended Data Fig. 3B. Neuropixels recorded spikes that were on average 5 times larger in amplitude than those recorded by Cambridge, yet the waveform shapes were similar. For example, both probes isolated MLI1 spikes with waveforms that exhibited a downward pattern in the molecular layer but an upward pattern near the Purkinje layer, consistent with a pinceau. We peak normalized each waveform and used the 2-D information (space across the probe, and time) to illustrate the SS, CS, and MLI1 spikes using UMAP. We found that SS, CS, and MLI1 waveforms clustered together across the two probes (Extended Data Fig. 3C). Thus, other than amplitude differences between Neuropixels and Cambridge, the spike waveforms for these two probes were similar in terms of their spatial pattern across the contacts, and in terms of their pattern across time.

We next examined spike interactions within cell types and between cell types, comparing Neuropixels and Cambridge (Extended Data Fig. 3D). Jitter corrected interactions for SS | SS, and MLI1 | MLI1 were very similar for the two probes. The ephaptic, GABA, rebound pattern of inhibition in SS | MLI1 were also quite similar in the two recordings. However, MLI1|MLI2, MLI1|CS, and MLI2|CS interactions were stronger in the Neuropixels recordings in the sense that within clique interactions were clearly greater than between cliques. Thus, it was easier to identify cliques with Neuropixels than Cambridge. This result highlighted the importance of using a probe that spanned many P-cell layers, allowing for sampling of multiple cliques. As a result, the larger span of Neuropixels better dissociated the between clique interactions from the within clique interactions.

*Computing the potent vector for each P-cell*

In this region of the cerebellum, the climbing fibers report to the cerebellum the direction of visuomotor events [90,91], likely because of the superior colliculus projections to the inferior olive [92]. For example, the climbing fibers reported to the cerebellum both the direction of the visual event, and then independently, the direction of the planned saccade [28]. We used the properties of the climbing fiber input to assign a potent vector to all the neurons that were members of the clique. The procedures were identical to those described in [10]. Briefly, we quantified this encoding by measuring the CS rate as a function of the direction of the visuomotor event during two different windows: 40 to 85 ms with respect to the visual cue onset, and -70 to +30 ms window from saccade onset. We then fitted a Von Mises distribution to the resulting rates as a function of angle [93]. This produced two vectors, one that quantified the strength and direction of the CS response to the visual events, and a second that quantified the strength and direction of the CS response to the motor events. Each vector had an angle $\theta$ and amplitude $\rho$, where the amplitude identified the sharpness of tuning. Thus, the amplitude of the vector was bounded by 0 (not tuned), and 1 (maximum sharpness). This produced two distinct vectors for each P-cell: one describing the tuning with respect to visual events, and one describing the tuning with respect to motor events. However, encoding of these two vectors were very similar: the magnitude of the vector for the visual event was highly correlated with the magnitude of the vector for the motor event. Hence, for each P-cell in each clique, we used the vector with the larger amplitude and labeled it as its *potent vector*. The direction of this vector defined $\theta$ in the various plots, for example, Figs. 3a & 3c.

*Computing the potent vector for each clique*

We organized the P-cells and the neighboring MLIs into neuronal cliques and then assigned a single potent vector to all the neurons in that clique. To do this, we considered cliques that had at least one P-cell (both SS and CS). We observed that the climbing fibers that projected to the same clique carried information that was much more similar to each other than between cliques [10]. To compute the potent vector for a given clique, we performed a vector average of the potent vectors of the P-cells that belonged to that clique, then assigned that single potent vector to all the P-cells and interneurons in that clique.

*Quantifying sub-millisecond spike interactions*

To compute a conditional probability, $\Pr(x(t+\Delta)=1|y(t)=1)$, that is, the probability of a spike in neuron $x$ at time delay $\Delta$, given that neuron $y$ produced a spike at time zero, we used a bin size of $w=0.1$ms and then aligned the data to each spike of neuron $y$ and computed the probability of observing a spike in neuron $x$ during the time period $\Delta \pm w/2$ . We then multiplied this probability by 10,000 and labeled that as Hz, i.e., spike rate per second. Next, we varied the time delay $\Delta$ and recomputed the probability for a new delay. Finally, to correct these probabilities for chance, we subtracted the jittered version of the same probability. For example, consider the conditional probability of a P-cell firing a simple spike, given that an MLI1 produced a spike at time zero (Fig. 1 e, SS | MLI1). If the two neurons are in a clique, immediately before the MLI1 spike there is a slight rise in the firing rate of the P-cell, which may be interpreted as common parallel fiber inputs to both neurons. Following the MLI1 spike, there is a sharp reduction in the firing rate of the P-cell: at 0.4ms delay, there is a 10Hz reduction in the simple spike rate. We interpret this as inhibition due to electrical interactions between the MLI and the P-cell at the pinceau. This inhibition is followed by a second inhibitory event at 1.3ms delay, which appears consistent with release of GABA from the MLI onto the P-cell. Finally, at a delay of 4ms there is a rebound in the probability of P-cell simple spikes, rising above zero.

*Quantifying joint probability of spiking in two neurons*

Our objective was to find a robust way to compute the probability of synchronous spiking with respect to chance among pairs of neurons, despite changes in their instantaneous firing rates.

The joint probability of spiking between two neurons, with spike times $a=\{a_1,a_2,\ldots,a_n\}$ and $b=\{b_1,b_2,\ldots,b_m\}$, for a given delay $\Delta$ and interaction window size w, is defined as:

$$\Pr(a_\Delta,b)=\Pr(|(a+\Delta)-b|\leq w/2) \tag{2}$$

Using Bayes theorem, we express the conditional probability (i.e., cross correlogram) as:

$$\Pr(a_\Delta|b)=\frac{\Pr(a_\Delta,b)}{\Pr(b)} \tag{3}$$

In practice, this formulation is equivalent to counting spike coincidences between the shifted spike train $a_\Delta$ and the spike times of neuron $b$ within the defined interaction window $w$. Analogous to firing rate estimation, we define a new spike train $c=\{c_1,c_2,\ldots,c_p\}$, consisting of the simultaneous spikes detected within window $w$ between spike trains $a_\Delta$ and $b$. Then, to assess whether observed synchrony reflects meaningful interactions beyond chance coincidences from firing rates, we employed interval jittering [26] to construct a null hypothesis of independent temporal structure. Specifically, the jittered spike train $b_{jitt}$ is generated as:

$$b_{jitt}=\delta\left\lfloor\frac{b}{\delta}\right\rfloor+\epsilon \tag{4}$$

where $\lfloor . \rfloor$ denotes the floor operator, and $\epsilon$ is a random uniform variable on $[0,\delta]$, where $\delta$ is the size of the jitter window. The synchrony hypothesis is tested by evaluating the following inequality:

$$\Pr(a_\Delta,b)\stackrel{?}{=}\Pr(a_\Delta,b_{jitt}) \tag{5}$$

Various statistical measures can quantify whether the measured synchrony is different than chance. Two widely used metrics are the synchrony index $\lambda$ and covariance:

$$\lambda_\Delta = \frac{\Pr(a_\Delta, b)}{\Pr(a_\Delta, b_{jitt})} \tag{6}$$

$$cov = \Pr(a_\Delta, b) - \Pr(a_\Delta, b_{jitt}) \tag{7}$$

Both measures capture excess spike coincidences but differ in interpretation: SI is a multiplicative ratio (division), reflecting the fold-change of synchrony relative to chance, whereas $cov$ reflects an absolute difference (subtraction) between observed and expected coincidence counts. Importantly, covariance also carries a biophysical interpretation, it quantifies the number of joint spikes beyond chance that could influence the downstream targets, providing a more direct link to potential synaptic impact.

However, as pointed out by Herzfeld et al. [8], it is critical to handle numerical instabilities in synchrony index calculation, especially in cases where $\Pr(a_\Delta, b_{jitt})$ approaches zero, such as during prolonged pauses in neuronal activity. To mitigate artifacts arising from divisions by near-zero values, we explicitly assign $\lambda_\Delta = 1$ whenever both $\Pr(a_\Delta, b)$ and $\Pr(a_\Delta, b_{jitt})$ are zero. This prevents artificial inflation of synchrony estimates, a misestimation that has been previously observed in spike synchrony analyses [9].

To robustly estimate $\Pr(a_\Delta, b_{jitt})$, we applied a bootstrapping procedure, performing 10 jitter iterations and averaging the results to obtain confidence intervals and reduce estimation variance.

*The joint-jitter plots*

To measure the empirical relationship between observed synchrony and chance synchrony, we plotted $\Pr(a, b)$ against $\Pr(a, b_{jitt})$ for each neuron pair during the entire recording session (Fig. 2c). To make this plot, for the neuron pair *a* and *b* we defined a new spike train $c = \{c_1, c_2, \ldots, c_p\}$, consisting of the simultaneous spikes detected during window $w = 1$ms. We then jittered one of the neurons and defined a new spike train $c_{jitt}$.

Next, to examine rate-dependent changes in synchrony, we divided each recording into non-overlapping 10 second estimation windows, independent of the animal's behavior. Within each window, we counted the number of joint spikes in the actual data, which provided an estimate of $\Pr(a, b)$, and compared it to the mean joint spikes after jittering one neuron's spike times, bootstrapped 10 times to estimate $\Pr(a_\Delta, b_{jitt})$. Each point in Fig. 2c represents a 10-second window, plotting the observed joint spikes against the expected joint spikes from jittering.

As the firing rates of the two neurons changed, so did the joint probability. By plotting the measured joint probability as a function of the joint probability of the jittered data, we produced the joint-jitter plots (Fig. 2c). The unity line in these plots represents the expected synchrony under the null hypothesis, i.e., independence. Points above this line indicate excess synchrony, while points below suggest mutual suppression or anti-synchrony.

By plotting the data for each cell pair in the joint-jitter format, we discovered that joint synchrony was approximately a linear function of jittered synchrony (Eq. 1). This implies that regardless of firing rates of the two neurons, the synchrony index $\lambda$ remains constant. The linear relationship between joint rates and jittered rates implied that $cov$ is not constant but varied with the firing rates. Thus, the joint-jitter plots demonstrated that synchrony index was a rate independent measure of synchrony, but not $cov$. We confirmed this prediction during saccades (Fig. 3), finding that SI remained constant.

For calculating the SI and cov during saccades we used similar joint spike train $c$ and $c_{jitt}$. We then aligned them to saccade peak velocity times and calculated the trial averaged joint and jittered responses. In order to reduce the noise, we smoothed each rate with a 10ms window moving average smoothing window. Covariance and SI were calculated by subtracting and dividing each time point respectively. Both joint-jittered plots, covariance, and synchrony index plots supported the constant linear relationship (Fig. 3) which matched the slopes found both from the whole trace normalized cross correlograms as well as whole session joint-jitter plots (Extended Data Fig. 6).

*Setting the width of the jitter window*

We employed interval jittering to construct a spike train that reflected the null hypothesis that the spike timing in the two neurons was independent of each other. To generate the jittered spikes, we used Eq. (4) to find the jittered spike times from the original spike times. An alternate method is spike-centered jittering where the spike is moved within a window centered at the location of the original spike. Prior work [27] has demonstrated that spike-centered jittering may be biased toward finding temporal structure where there is none (i.e., false positives). In contrast, interval jittering appears to not have this issue. Thus, here we employed the interval jittering technique.

A critical decision was regarding the size of the jittering interval, i.e., $\delta$. The value of $\delta$ is required to be large enough so that it keeps the single spike effect intact, while at the same time small enough so that it does not perturb the average firing rate dynamics of the cells. Given the fast nature of saccades and fast dynamics of the pMLI1s our choice of $\delta$ was more restricted than common 20ms for cerebral cortex. However, the submillisecond ultra-fast nature of fast interactions between P-cells and MLI1s allowed us to lower $\delta$ to 5ms. To test the effect of $\delta$ on our results, we tested a range of values from 3ms to 100ms (Extended Data Fig. 10B). As P-cells have slower rate dynamics and lower rate correlation P-cell cross correlograms were highly robust to the choice of $\delta$ (Extended Data Fig. 10B). However, the pMLI1s showed a substantial rate correlation which could be observed both on the jittered cross-correlograms as well as in between-clique interactions. This is consistent with the homogeneous fast dynamics of the pMLI1s with average baseline activities of ~30Hz reaching to 90Hz peaks in 10s of milliseconds for all directions of saccades. Due to fast nature of pMLI1 rates we picked $\delta$=5ms to prevent overestimation of observed spike synchrony due to the rate correlations.

*Measuring synchrony during saccades*

We began by transforming the synchronous spike events across the entire recording into a new spike train, $c = \{c_1, c_2, \ldots, c_p\}$, representing joint spikes between neurons *a* and *b*. We then generated 10 surrogate spike trains by jittering. Next, we aligned these joint spikes to the peak velocity of saccades and computed peri-stimulus time histograms (PSTHs) to estimate $\Pr(t, a, b)$ and $\Pr(t, a, b_{jitt})$, where $t$ denotes time relative to saccade peak velocity. We smoothed both joint and jittered probabilities using a 10ms moving average window. Using these two smoothed time-dependent probabilities, we computed synchrony index and covariance as a function of time (Eq. 6 & 7). We fixed the numerical instability of division in SI through 10ms smoothing of joint and jittered traces in time and averaging 10 bootstrapping samples to prevent divisions by zero. Finally for all SI calculations, whether whole recording or aligned to the saccades, we accepted the null hypothesis SI=1 when the jittered values were zero.

*Testing the validity of the spike interaction measures*

We measured joint spike probabilities in three different ways. First, we calculated $\Pr(a_{\Delta}, b)$ and $\Pr(a_{\Delta}, b_{jitt})$ across the entire recording session for time delays ($\Delta$) ranging from –10 ms to +10 ms, using a fine-grained coincidence window of $w = 0.1$ms. This method produced results similar to cross-correlograms but normalized by the firing rates (Fig. 2b). However, while informative, this approach only provided a time-averaged picture and did not reveal how the joint probability varied as a function of the rates of the two neurons.

Second, to examine rate-dependent changes in synchrony, we divided each recording session into non-overlapping 10-second estimation windows, independent of the animal's behavior. Within each window, we counted the number of joint spikes $\Pr(a_{\Delta}, b)$ in the actual data and compared it to the mean joint spikes after jittering one neuron's spike times, bootstrapped 10 times to estimate $\Pr(a_{\Delta}, b_{jitt})$. This produced the joint-jitter plots. Remarkably, these plots revealed a linear relationship: as the jittered joint spiking rate increased, the observed joint spiking also increased linearly, consistently diverging above the unity line. This indicated that as cell pairs increased their firing rates, they also aligned their spikes more precisely than expected by chance, resulting in a rate-dependent covariance. Furthermore, different neuron pairs exhibited distinct slopes, allowing us to classify them into groups with varying synchrony indices (Fig. 2f). We interpreted these slopes as biophysical manifolds that characterized how synchronous spike probability scaled with joint firing rates for each cell pair. This is because the x-axis of these plots is related to the firing rates of each neuron as follows:

$$\Pr(a, b_{jitt}) \approx \Pr(a)\Pr(b) \quad (8)$$

Third, to test whether these synchrony manifolds changed during behavior, specifically during saccades, we transformed the synchronous spike events into a new spike train, $c = \{c_1, c_2, \dots, c_p\}$, representing joint spikes. We also generated 10 surrogate spike trains via jittering for comparison. We then aligned these joint spikes to the peak velocity of saccades and computed peri-stimulus time histograms (PSTHs) to estimate $\Pr(t, a, b)$ and $\Pr(t, a, b_{jitt})$, where $t$ denotes time relative to saccade peak velocity. We smoothed joint and jittered probabilities in time using a 10ms moving average window. Plotting $\Pr(t, a, b)$ against $\Pr(t, a, b_{jitt})$ during the saccade epochs revealed that the synchrony remained confined to the same biophysical manifold identified in the second approach, maintaining a consistent slope throughout the movement (Fig. 3c). This result suggested that the spike-time coordination between MLI1s and P-cells was not dynamically modulated by the behavior itself but was a stable, rate-dependent property of the circuit that scaled with activity levels.

*Measuring interactions between MLIs and P-cells via conditional probabilities*

The corrected joint probability, i.e., $\Pr(a, b) - \Pr(a, b_{jitt})$ measures how many additional spikes per unit of time are being conveyed to downstream targets by a pair of neurons, beyond what would be expected by chance. This makes joint probability a meaningful metric for inferring the graphical structure of the circuit, particularly when assessing whether spike alignments from a neuron pair could influence a common downstream target. However, to evaluate the effect of one neuron on another, such as the effect of MLI1s projecting onto P-cells, the conditional probability is a more appropriate measure.

The conditional probability $\Pr(a_{\Delta}|b)$ quantifies how the firing rate of a postsynaptic neuron $a$ changes aligned to the spike times of a presynaptic neuron $b$, with a time delay ($\Delta$) parameter explicitly capturing how this interaction evolves across different temporal offsets. To determine whether the observed suppression reflects a true causal interaction rather than a byproduct of correlated firing rates, we extended our interval jittering approach to statistically assess the significance of these interactions. This method effectively decomposes rate correlations driven by common parallel fiber inputs, isolating fast, spike-induced suppressions that arise from direct inhibitory interactions.

Fig. 4a illustrates a spectrum of P-cell suppression profiles following MLI1 spikes. While some P-cells exhibited only a sharp, fast-latency suppression consistent with pinceau-mediated ephaptic inhibition, others display a secondary, more prolonged suppression, reflecting an increasing contribution of GABAergic synaptic inhibition. The larger the GABA component, the more pronounced the post-suppression rebound, revealing a gradient of inhibitory influence across different P-cell–MLI1 pairs.

A key concern in interpreting suppression based on spike probabilities is that we are not directly measuring the inhibitory currents, but rather the probability of spike omissions. This introduces a limitation: if the postsynaptic P-cell is silent at a given moment, an inhibitory input cannot further suppress its firing probability. Consequently, our conditional probability measure may underestimate inhibition effects at low firing rates. This limitation arises from the measurement method, not from a failure of the inhibitory mechanism itself.

To mitigate this, we introduced a firing-rate-normalized conditional probability measure that conditioned the observed suppression not only on the presynaptic spike timing but also on the instantaneous firing rate (IFR) of the postsynaptic neuron (Fig. 4a). This method built upon prior work on 3D auto-correlograms [8], where autocorrelation structures of single neurons were conditioned on their own instantaneous firing rates. Our extension generalized this concept to cross-correlograms between two neurons by conditioning the postsynaptic neuron's response to presynaptic spikes based on its instantaneous firing rate dynamics at the time of the interaction. We defined this conditional probability as $\Pr(a_{t+\Delta}|b_t, r_a(t))$, where $r_a(t)$ is a smoothed estimate of the instantaneous firing rate of neuron $a$ at the time of each presynaptic $b$ spike.

For each spike of neuron $b$, we assigned the corresponding instantaneous firing rate of neuron $a$ at that moment. We then compute the conditional probability by considering all spikes of neuron $a$, but only a subgroup of neuron $b$ spikes that occur when neuron $a$'s firing rate fell within a specified range. This allowed us to analyze how postsynaptic response varied as a function of local firing rates (Fig. 4a). This approach allowed for fair comparisons across neuron pairs with differing baseline firing rates, ensuring that the suppression of the MLI on the P-cell was not confounded by periods of inactivity in the P-cell.

*Effect of the synchronous MLI1 spikes on the downstream P-cell*

We investigated whether synchronous spikes from multiple MLI1s could cumulatively enhance the suppression of a downstream P-cell (Fig. 4b). Given the rapid and potent inhibitory effect of a single pMLI1 spike on a P-cell, it was plausible that synchrony within the upstream pMLI1 population played a functional role by summing their effects (superposition) to induce a stronger suppression. To test this, we identified triplets of neurons consisting of two connected pMLI1s and their common downstream P-cell and computed the following conditional probability: $\Pr(ss_{t+d}|a_t, b_{t-\Delta}, r_{ss}(t))$. In this expression, $a_t$ is MLIa spiking at time t, $b_{t-\Delta}$ is MLIb spiking at some time earlier, $r_{ss}(t)$ is the instantaneous firing rate of the P-cell at time t (see below for more detail on instantaneous firing rate calculation), and $ss_{t+d}$ is the probability of the simple spike occurring at a time after the MLIa spike. As shown in Fig. 4b, sequential pMLI1 spikes evoke largely independent suppressive effects on the P-cell when separated by a couple of milliseconds. However, when the two pMLI1 spikes occurred within a narrow time window (within a millisecond), their suppressive effects added, resulting in a significantly larger suppression of the P-cell's firing probability. This observation suggests that spike synchrony within the pMLI1 population can effectively modulate the strength of inhibition exerted on P-cells through superposition.

We further tested the superposition of two pMLI1s in two ways. If the principle of superposition holds, the combined effect of two MLI spikes with a temporal delay should be predictable by adding their individual, time-shifted suppression profiles on the downstream P-cell. We evaluated this by comparing the observed and estimated suppression across three temporal phases: the pinceau period (0–0.8 ms after the MLI spike), the GABAergic period (0.8–2.5 ms), and the rebound period (2.5–6 ms), as shown in Fig. 4c (dashed lines: expected; solid lines: observed). To quantify uncertainty of estimation, through bootstrapping (n=1000), we computed 95% lower and upper confidence intervals by jittering the timing of one MLI within a 1 ms window, matching the delay range between the two MLIs.

*Computing instantaneous firing rates*

To compute the instantaneous firing rate $r(t_s)$ of a neuron, where $t_s$ is the time that a spike occurred, we used an adaptive smoothing window. First, the raw instantaneous firing rate was defined at each time point $t_s$ as the inverse of the inter-spike interval between the two spikes immediately preceding and following $t_s$. Next, we used an adaptive algorithm to compute a smoothing window size for each $t_s$. The width of this smoothing window was 700 ms divided by the number of spikes contained in a 100 ms window centered on $t_s$. These parameters were chosen so that, on average, the smoothing window contained approximately 7 spikes. The instantaneous firing rate, $r(t_s)$ at each spike time $t_s$ was the average of the raw firing rate within a window centered on $t_s$, with duration equal to the smoothing window duration described above. Simulations were used to determine the window size needed to prevent overfitting the firing rate to the spike times.

To test whether our algorithm for estimating the instantaneous firing rate was accurate, we performed simulations via a P-cell model (see below) for which the ground truth regarding firing rate was known. In these simulations (Extended Data Fig. 10) we confirmed that our algorithm for estimating the neuron's instantaneous firing rates was accurate.

*Phase normalization of spike timing*

Consider a neuron whose spike timing remains regular despite changes in firing rates. To understand spike timing of this neuron, imagine a clock that runs from 0 to $2\pi$ at a speed that is defined by the instantaneous firing rate of that neuron. Thus, as the firing rate increases, the clock runs faster. This neuron can generate a spike at any time beyond its refractory period, but because the neuron is regular, if it has generated a spike at time 0 then the timing of the next spike will be a probability distribution that has a strong peak centered around $2\pi$, no matter what its instantaneous firing rate may be. If the neuron is not very regular, the spike timing will be widely distributed with a weaker peak around $2\pi$.

For example, the P-cells in Fig. 6b have spike timing with an initial peak at $2\pi$, followed by a second peak at $4\pi$. In contrast, the pMLI1s have a more uniform distribution. Thus, the P-cells have a greater reliance on an internal clock than the pMLI1s.

Using the internal clock analogy, we set out to measure the spiking probability for each neuron in terms of the phase of its internal clock, where the speed with which the internal clock runs is defined by that neuron's instantaneous firing rate $r(t_s)$, where the time of a spike is specified by $t_s$. In other words, the phase was defined locally at each spike time.

Given that the neuron spiked at time $t_s$, the phase $\vartheta$ with respect to $t_s$ at time $t$ was defined as:

$$\vartheta(t|t_s) = 2\pi\ (t - t_s)\, r(t_s) \tag{9}$$

In this equation, the time difference $t - t_s$ is measured in seconds, while the firing rate is $r(t_s)$ is measured in spikes/sec. Thus, their product is a unitless quantity that is equal to one when the time elapsed equals the inverse of the instantaneous firing rate. The product $(t - t_s)\, r(t_s)$ represents the ratio of the time elapsed to the expected time of the next spike. We then scale by a factor of $2\pi$ to express this ratio in the form of the phase of an oscillator in units of radians, such that it completes one full cycle (i.e. $2\pi$ radians) within the expected interspike interval.

The conditional probability plot in Fig. 6c (left subplot), where firing rate is represented as spikes/cycle, was constructed by iterating over each spike time $t_s$ in the spike train and obtaining the phase $\vartheta(\ t_s' |\ t_s)$of every other spike time $t_s'$ with respect to the current spike, then plotting a histogram of the frequency of occurrence of each phase. We used a bin size of $2\pi/50$ radians, or 1/50 of a full cycle of the neuron's internal clock. The probability of a spike occurring in each phase bin was normalized by dividing by the bin size to obtain a firing rate in units of spikes/cycle. This is analogous to how traditional time-based spiking probabilities are normalized to spikes/second.

To give an intuition regarding Fig. 6c, consider the following. By definition, a neuron fires on average one spike per cycle, since 1 cycle is the length of its expected inter-spike interval, so the phase probability plot for a pure Poisson neuron with independent spike times and no refractory period would be a flat line with a value of 1. If the normalized spiking probability for a neuron at a given phase $\vartheta$ is greater than 1, then the spike times tend to be concentrated at that part of the cycle, whereas if the normalized spiking probability is less than 1, then the spikes are less likely to occur at that part of the cycle than they would be for a Poisson neuron of the same rate.

*Phase resetting via ephaptic interactions*

Having established a method for estimating the phase of a neuron's internal clock, we next examined how the phase of the internal clock affected interactions with neighboring cells. For each within-clique pair of P-cells, we found the conditional probability of PCa spiking as a function of the timing of a PCb spike relative to PCa's clock. Specifically, we binned pairs of PCa and PCb spikes by the phase of PCa's internal clock when the PCb spike occurred, $\vartheta(t_b\ |\ t_a)$. Again, we used a bin size of $2\pi/50$ radians, such that a pair of spikes $(t_a, t_b)$ fell within a particular bin centered at $c$ if $|\vartheta(t_b \mid t_a) - c| < \pi/50$. Next, we iterated over all pairs $(t_a, t_b)$ within each bin and found the phase $\vartheta(t_a{}' \mid t_a)$ of every other PCa spike with respect to $t_a$, and then obtained a histogram of the frequency of occurrence of each phase, as described above. This gave us the spiking probability of PCa as a function of its clock phase conditioned on a PCa-PCb interval, $\vartheta(t_b \mid t_a)$. Representative examples for three different values of $\vartheta(t_b \mid t_a)$ are shown in Fig. 6c (top). These plots reveal how the effectiveness of ephaptic coupling for inducing spikes in a given cell is modulated by that cell's clock phase. As a control, we did the same analysis after jittering the spike train of PCb, finding that the change in spiking probability was dependent on precise spike timing.

Next, we asked whether an ephaptic interaction with a neighboring cell could reset the phase of a P-cell's internal clock. For instance, if PCb induced an "off-cycle" ephaptic spike in PCa, such as when PCa's phase was at $\pi$, would the next PCa spike also be off-cycle, or would its phase return to what it would have been in the absence of an ephaptic spike? In other words, did an ephaptic interaction affect the behavior of a P-cell only transiently or did it induce a long-term change in the cell's spike timing? To answer this question, we again considered within-clique pairs of P-cells, PCa and PCb, and classified pairs of spike times $(t_a, t_b)$ as putative ephaptic spiking events if $t_a$ occurred less than 0.5 ms after $t_b$. We then repeated the analysis described above, binning each pair $(t_a, t_b)$ by the normalized interspike interval $\vartheta(t_b \mid t_a)$ and finding a histogram of PCa interspike intervals $\vartheta(t_a{}' \mid t_a)$ for each bin, but this time we restricted our analysis to only include pairs $(t_a, t_b)$ where $t_b$ was part of a putative ephaptic event. We also restricted PCa spike times $t_a{}'$ to those that occurred after the putative ephaptic event, because we were interested in understanding how the ephaptic event affected the P-cell's spike timing going forward. Results for the same three values of $\vartheta(t_b \mid t_a)$ are shown in Fig. 6c (bottom row). The control is the same as in Fig. 6c (top row).

To summarize the results from Fig. 6c, we plotted the probability that a PCb spike induced an ephaptic spike in PCa as a function of PCa's phase (Fig. 6d, left). To do this, we iterated over each bin of normalized PCa-PCb intervals, $\vartheta(t_b \mid t_a)$, and then counted the number of putative ephaptic spiking events divided by the total number of spike timing pairs $(t_a, t_b)$ in each bin. Some of these putative spiking events will occur purely by chance, rather than being the result of an ephaptic interaction. Therefore, to estimate the true ephaptic spiking probability, we repeated the same analysis after jittering the spike times of PCb and subtracted the jittered probability from the true probability.

Fig. 6c (bottom) shows that an ephaptic spike at off phase (e.g. at $\pi$ or $3\pi$) induced a phase change in the cyclical spike timing probability of the P-cell, while an ephaptic spike induced on phase (at $2\pi$), produced less of a phase change. To quantify this phase change, we found the expected change in PCa phase after a putative ephaptic event as a function of the phase of PCa at the time of the ephaptic event. As above, we found all spike timing pairs $(t_a, t_b)$ where $t_b$ was part of a putative ephaptic event and binned them by the normalized PCa-PCb interval $\vartheta(t_b \mid t_a)$. For each spike pair $(t_a, t_b)$, we found the phase of the first spike after the putative ephaptic event, $\vartheta(t_a{}' \mid t_a)$. This gives us a distribution of phases which we wish to compare to a control that shows what the distribution would have been had there not been a putative ephaptic event. The first PCa spike after a putative ephaptic event is of course the second PCa spike after $t_b$ (since the first one is included in the ephaptic event), so a useful control is the distribution of the phase of the second PCa spike after a spike in the jittered PCb train, again binned by $\vartheta(t_{b,jitt} \mid t_a)$.

Phases are circular values, so the expected phase change is not simply the difference between the means of these two distributions (because a phase change of $2\pi$ is equivalent to no change at all). Instead, we found the difference between the circular means of each distribution. Let $\vartheta_k^{reset}$ denote the k-th instance of $\vartheta(t_a' \mid t_a)$ following an ephaptic spiking event for a given PCa-PCb interval bin, and let $\vartheta_k^{control}$ be the k-th instance of $\vartheta(t_a' \mid t_a)$ following a jittered PCb spike. Then the angle difference between the two distributions is $\Delta\vartheta = \arg(z)$, where

$$z = \sum_k \exp(i * \vartheta_k^{reset}) \sum_k \exp(-i * \vartheta_k^{control}) \quad (10)$$

To average over all P-cell pairs, we found the complex vector z corresponding to each pair and then defined the mean expected phase change as $E[\Delta\vartheta] = \arg(\sum_i z_i)$. For the sake of clarity in the plots, phase shifts near +/-π were assigned to be either positive or negative based on the surrounding values, preventing any abrupt discontinuities resulting from wrapping a circular values onto a linear graph. 95% confidence intervals were determined by bootstrapping.

*Modeling ephaptic coupling among P-cells*

To investigate the role of ephaptic coupling in coordinating spike timing among P-cells, we developed a two-compartment linear integrate and fire (LIF) model (soma and axon initial segment, AIS) of pairs of P-cells where somatic membrane potential evolved according to traditional LIF [5,17,21].

$$c_m \frac{dV_m}{dt} = \Sigma I \quad (11)$$

In above, $c_m$ represents membrane capacitance and $V_m$ represents membrane voltage. At each time step, the change in P-cell somatic membrane voltage $V_m$ was determined from the net sum of various input currents: a leak current $I_L$, a net input current $I_{net}$ mapped from a target rate function $r_i(t)$, and a net membrane noise $\sigma$:

$$c_m \frac{dV_m}{dt} = (-I_L + I_{net} + \sigma) \quad (12)$$

The leak current at each time point was calculated as the driving force of the membrane voltage at the prior time step scaled by the leak conductance. Here, $E_L$ is the membrane reversal potential (-70 mV) and $g_L$ is the leak conductance (0.5 $\mu S$):

$$I_L = g_L(V_m(t-1) - E_L) \quad (13)$$

The net current at each time point was calculated via a target instantaneous rate $r_i(t)$ scaled linearly by a constant $\alpha$. This was numerically set to produce the target firing rates given constant membrane leak conductance $g_L$ and capacitance $c_m$ values. The target rate function was built from saccade-aligned simple spike rates from actual P-cells (dataset size *n* = 226, with *n* = 101 bursters and *n* = 125 pausers). For each simulation, a pair of P-cells (burster-burster, burster-pauser, or pauser-pauser) was chosen, and two matching real P-cell profiles were randomly selected from the dataset, interpolated from 1 kHz to 30 kHz to match the model timestep, and tiled across trials to span multiple behavioral periods:

$$I_{net,i} = \alpha r_i(t) \quad (14)$$

The net noise term $\sigma$ was a zero mean Gaussian noise with scaling factor $I_\sigma$. Here, $I_\sigma$ controlled spike-train regularity: a low $I_\sigma$ value represented little random noise input to the P-cell, and therefore high regularity, whereas a high $I_\sigma$ value represented large random noise input to the P-cell, and therefore low regularity with non-significant effect on average firing rates:

$$\sigma = I_\sigma \, \xi(t) \qquad \xi(t) \sim \mathcal{N}(0,1) \tag{15}$$

Once the net current at a given time point was determined, the soma membrane voltage of each P-cell was updated using forward Euler integration.

Each neuron's AIS voltage $V_{AIS,i}$ is the sum of two components: an intrinsic voltage component which follows that neuron's somatic voltage $V_{m,i}$, and a weighted voltage $V_{ex,j}$ which is the voltage perturbation driven by the extracellular field produced by spikes in the nearby neurons through coupling matrix $A$:

$$V_{AIS,i}(t) = V_{m,i}(\mathrm{t}) - \sum_{\mathrm{j}} \mathrm{A_{ij}} \, V_{ex,j}(t) \tag{16}$$

Here, $A_{ij}$ is the i-th row and j-th column of an nxn connectivity matrix. For our simulation we consider $A$ as a symmetric 2x2 coupling matrix which was determined by the AIS-AIS distance $d$ ($\mu m$) between the simulated pairs of neurons using the distance scaling rule from [5] where $k_0$ is a geometry/offset constant that captures out-of-plane separation between the neurons and avoids numerical issues at $d = 0$:

$$A = \begin{bmatrix} 0 & h(d) \\ h(d) & 0 \end{bmatrix} \quad h(d) = \frac{k_0}{d^2 + k_0} \tag{17}$$

Each neuron's extracellular field $V_{ex,j}(t)$ is the convolution of its spike train $s_j(t)$ with an extracellular kernel $k_{Ve}(t)$ constructed from the AIS spike waveform (Eq. 18). When modeling ephaptic inhibition at the MLI1-PC pinceau, Blot and Barbour [21] approximate the extracellular voltage at the pinceau as proportional to the negative derivative of the intra-pinceau potential. We apply this same approximation for our extracellular kernel and model the extracellular voltage as proportional to the negative derivative of the intrinsic voltage at the AIS. For ephaptic coupling, the relevant intrinsic voltage at the AIS is the extracellular simple spike waveform, which we model using a gamma kernel $k_\gamma(t, \alpha, \theta)$.

$$s_j(t) = \sum_k \delta\left(t - t_{k,j}\right) \tag{18}$$

$$k_{Ve}(t) \propto -\frac{d}{dt} k_\gamma(t, \alpha, \theta)$$

$$V_{ex,j}(t) = \left(s_j(t) * k_{Ve}(t)\right)$$

A neuron spikes when the total voltage at the AIS exceeds the threshold voltage (-55 mV) at a given time point, and the neuron is not already in a refractory period. Upon a spike, $V_m$ and $V_{AIS}$ are both clamped to $E_L$ for a 1 ms refractory period during which no new spikes can be emitted. Supplementary Table 1 provides the values for the parameters used in the P-cell simulations. We matched the parameters of the kernel to get a cross-correlogram shape that resembled real cross-correlograms in data (Figs. 1e, 5d).

*Modeling gap-junction among pairs of MLI1s*

To investigate the role of gap junctions in coordinating spike timing in MLI1s, we developed a conductance-based leaky integrate-and-fire model (Eq. 19):

$$c_m \frac{dV}{dt} = \Sigma I \tag{19}$$

At each time step, the change in model MLI1 membrane voltage was determined from the net sum of various input currents: a leak current $I_L$, an excitatory current derived from simulated parallel fiber (PF) input $I_{ex}$, a gap junction current $I_{gap}$, and a bias current $I_b$. Once the net current at that time point was determined, the membrane voltage was updated using forward Euler integration.

$$\frac{dV}{dt} = \frac{1}{C_m}\left(-I_L - I_{ex} - I_{gap} + I_b\right) \tag{20}$$

The leak current at each time point was calculated as the driving force of the membrane voltage at the prior time step scaled by the leak conductance (Eq. 21). Here, $E_L$ is the membrane reversal potential (-70 mV) and $g_L$ is the membrane leak conductance (2.5 nS).

$$I_L = g_L(V_m(t-1) - E_L) \tag{21}$$

The excitatory current at each time point was calculated as the glutamatergic driving force scaled by an excitatory conductance kernel (Eq. 22). Here, $E_{ex}$ is the reversal potential of the excitatory glutamatergic PF-MLI synapse (0 mV), and $g_{ex}$ is a kernel basis function designed to mimic the time course of a glutamatergic synaptic transient:

$$I_{ex} = g_{ex}(V_m(t) - E_{ex}) \tag{22}$$

To properly mimic the time course of an AMPA excitatory postsynaptic current (EPSC) at the PF-MLI synapse, we followed standard convention for modeling AMPA synaptic transients in existing computational modeling literature and modeled the excitatory conductance kernel as a difference of exponentials parametrized by $t, \tau_1$ and $\tau_2$, where $t$ is the time in ms since PF spike arrival, and $\tau_1$ and $\tau_2$ are respectively the rise and decay time constant parameters of the AMPA synaptic transient. The values of $\tau_1$ = 0.166 ms and $\tau_2$ = 2.938 ms were determined sourcing a variety of AMPA synaptic transient time constant values from existing computational modeling or stimulation papers and then fitting a difference of exponentials function to the average AMPA model (Eq. 23) [94–98].

$$g_{ex} = g_{e_{max}} exp\left(-\frac{t}{\tau_2}\right) - exp\left(-\frac{t}{\tau_1}\right) \tag{23}$$

To determine the excitatory conductance at a given time point, each PF's spike train was convolved with the excitatory conductance kernel. Then, each PF's excitatory conductance value at that time was weighted by the randomly assigned synaptic strength of that given PF-MLI connection. The final excitatory conductance for that time point was then calculated as the sum of all weighted PF conductances at that time point.

The spike train for each PF was generated via a non-homogeneous Poisson process with a post-spike refractory period of 0.5ms. First, a candidate spike $t_i$ was drawn from a homogeneous Poisson process parameterized by a constant rate ($t_1, t_2, \cdots, t_N \sim Poisson(\lambda_{max})$). If the candidate spike occurs at least 0.5 ms after the most recent spike, it was then either accepted or rejected via a thinning process (a spike at time $t_i$ is accepted with probability $\lambda(t_i)/\lambda_{max}$). This made the Poisson rate function dynamic and thereby rendered the Poisson process non-homogeneous (Eq. 24).

$$\boldsymbol{Pr}(PF_n \text{ spikes at } t_i|t_{i-1}) = \frac{\lambda(t_i)}{\lambda_{max}} \cdot \mathbb{I}\left[t_i - t_{i-1} \geq t_{ref}\right] \tag{24}$$

This effectively ensured that the frequency of PF spikes over time corresponded to the input time-dependent instantaneous rate function $\lambda(t)$ which served as a proxy for behaviorally driven PF excitation.

Granule cells typically receive presynaptic inputs from 4 mossy fibers [99], which themselves receive input originating from all over the brainstem and cerebral cortex. In the oculomotor vermis of the cerebellum, "state" mossy fibers convey efference copies of a saccadic movement command with various delays to the downstream PFs [10,83,84,86]. In Eq. (24), $\lambda(t)$ is a sequence of evenly spaced kernel functions (3 per 1000 ms) designed to represent the average firing rate of saccade-responsive state mossy fibers.

To simulate the excitatory input that a given PF might receive, we used our recorded data from n = 222 "state" MFs [10]. For each PF, we randomly determined how many MF inputs it would receive (3-5) and then randomly selected that many MFs from the real data. Each selected MF input was assigned a random synaptic weight (0-1). The selected MF average responses during task-relevant saccades were then scaled according to the assigned synaptic weight, and the average of the weighted MF rates was then taken and normalized to serve as the basis function for that PF.

Additionally, each PF-MLI connection was assigned a random, nonnegative synaptic weight (0-1). Using these PF inputs, we simulated 3 saccades per second with a random scaling of peak velocity (uniform between 0 and 4). Assuming linear relationship between the rates and peak velocities [10,83,84,86], for each saccade we modeled the PF rate in time scaled according to the peak velocity to form $\lambda(t)$.

To avoid any edge artifacts at the end of the PF basis function, we interpolated from the last value of the basis function to the $\lambda(t)$ baseline value of 0 using MATLAB's implementation of the Piecewise Cubic Hermite Interpolating Polynomial algorithm [100].

The gap junction current at each time point was calculated as the membrane voltage of the paired MLIs at the prior time point, scaled by the gap junction conductance $g_{gap}$.

$$I_{i,j\ gap} = g_{i,j\ gap}\ (V_m^i(t-1) - V_m^j(t-1)) \tag{25}$$

Here, $V_m$ represents the voltage difference between the two MLIs and $I_{i,j\ gap}$ represents the current going from neuron j to neuron i. For networks with more than two MLIs, the gap junction current was additionally scaled by the Laplacian matrix $\mathcal{L}$.

$$I_{gap} = g_{gap} \cdot \mathcal{L}_{ij} \cdot V_m(t-1) \tag{26}$$

Here, the graph Laplacian represents the adjacency structure of the MLIs in the network and was computed from a randomly generated symmetric adjacency matrix ($A\_\{ij\} \in \{0,1\}$) with connection probability $p$.

$$\mathcal{L} = \text{diag}\left(\sum_j A_{ij}\right) - A \tag{27}$$

The stochastic bias current at each time point was comprised of a constant component ($I_c$ = 0.028) and a Gaussian white noise implemented using the Euler-Maruyama method with diffusion amplitude of $\sigma$ = 0.0158 nA $\sqrt{ms}$. This amplitude is equivalent to a Gaussian current-noise standard deviation of $I_\sigma$ = 0.05 at the reference timestep of $\Delta t_{ref}$= 0.1 ms. Together, these components generated the randomly fluctuating background input that gives the MLIs a baseline firing rate:

$$I_b = \frac{I_c}{C_m} + \frac{\sigma}{C_m}\sqrt{\Delta t\, \xi(t)} \qquad \xi(t) \sim \mathcal{N}(0,1) \tag{28}$$

Once the net current was calculated, the membrane voltage of each MLI was updated using the Forward Euler method. During the simulation, any neuron which had a membrane voltage above threshold entered a refractory period for 0.5 ms during which its voltage was clamped to a predefined spike shape. To investigate the role that gap junction conductance plays in shaping MLI synchronous spiking, we simulated a network with 10 PFs and 2 MLIs 120 times for four different gap junction conductance values. The parameter values for the MLI simulations are provided in Supplementary Table 2.

*Measuring the effect of synchrony on saccades*

To determine whether synchronous spiking had any measurable effects on saccades, we focused on primary saccades, then compared trials in which pairs of P-cells exhibited an unusually high number of synchronous spikes to trials in which those same pairs exhibited an unusually low number of synchronous spikes. To do this, we calculated the expected number of synchronous spikes for each P-cell pair by first jittering the spike trains and counting the average number of synchronous spikes in the jittered spike trains in the window from -20 ms before saccade peak velocity to +10 ms after peak velocity. We then multiplied this value by the synchrony index of the pair, determined using the spike trains over the entire recording. Since we show that true synchronous spiking rate is linearly related to the jittered synchronous spiking rate, with slope determined by the synchrony index, this method gives us an unbiased estimate of the expected number of synchronous spikes. However, because the temporal window is so brief, the expected number is usually close to 0, making comparisons with the real synchronous spike count noisy and unreliable. To mitigate this, we add up the expected number of spikes across all P-cell pairs in a clique, operating under the assumption that P-cell pairs in the same cliques should induce similar behavioral consequences. Only cliques with at least 4 P-cells were included. We then compared this number with the sum of the actual number of synchronous spikes across all pairs in the clique. An example from a single clique is shown in Extended Data Fig. 7A. Trials in which the actual number of synchronous spikes was higher than the expected number are classified as high synchrony.

To verify that this method reliably separates high and low synchrony trials, we plotted the average synchronous spiking rate across all pairs of P-cells that were included in the analysis for high synchrony trials and low synchrony trials, binned by saccade direction relative to the CS-on direction of the clique, $\theta$. The results show that synchronous spiking rates are clearly different on high and low synchrony trials (Extended Data Fig. 7B, top), while overall P-cell firing rates are not significantly different for high and low synchrony trials (Extended Data Fig. 7B, bottom), showing that the method is minimally confounded by P-cell rates.

Having identified high synchrony and low synchrony trials, we attempted to see if there were any behavioral differences between the cases. We hypothesized that variation in the amount of P-cell synchrony should affect behavior along the potent vector of the P-cells, labeled as $\theta$. To test this, for each clique, we found the difference between the average saccade aligned velocity vectors for the high synchrony trials and low synchrony trials, then projected this difference onto the potent vector of the clique. We binned results by saccade direction relative to $\theta$ and averaged over all cliques (Extended Data Fig. 7C). A 95% confidence interval (Extended Data Fig. 7C, gray) was found via permutation testing by shuffling the labels of the high synchrony and low synchrony trials. None of the directions showed a significant effect. Thus, we could not detect the effect of synchronous P-cell spiking on saccades.

*Statistics and Reproducibility*

We performed t-tests, or rank sum tests, or ANOVAs, to compare distributions. To test the effects of MLI1 spike timing on P-cell suppression and rebound (Fig. 4e), we leveraged the approximately linear relationship between suppression/rebound and instantaneous firing rate (IFR) within each inter-spike interval (ISI) condition. Because variability in triplet connectivity strength (2 pMLI1s converging on a P-cell) can influence the IFR interaction slope, we used a linear mixed-effects model to represent the data:

$$y_{i,k,r} = (\beta_k + b_i)r + \left(\beta_k^0 + b_i^0\right) + \epsilon \tag{29}$$

In the above equation, $y_{i,k,r}$ represents the suppression/rebound corrected conditional firing rate of the P-cell, $r$ is the IFR, $k$ is ISI bin, and $i$ is the neuron triplet number. $\beta_k$ is the slope of the relationship and $\beta_k^0$ is the intercept for each ISI bin (fixed effects), whereas $b_i$ and $b_i^0$ are the triplet specific deviations of slope and intercept (random effects). Thus, ISI is treated as a categori cal variable with four levels corresponding to pMLI1 pair conditions (0, 2, 4, and 6 ms). This model estimated ISI-specific fixed effect slopes $\beta_k$ for the relationship between P-cell firing rate and IFR, while accounting for the triplet specific variability in connection strength.

No statistical method was used to predetermine sample size. No data were excluded from the analyses. The experiments were not randomized. The Investigators were not blinded to allocation during experiments and outcome assessment.

**Data availability:** The data used to generate the figures are available at https://osf.io/fc7aw/overview?view_only=626276ef70d4451cab45be2a3418fa27

**Code availability**: All code used to generate the figures and simulations are available at https://osf.io/fc7aw/overview?view_only=626276ef70d4451cab45be2a3418fa27

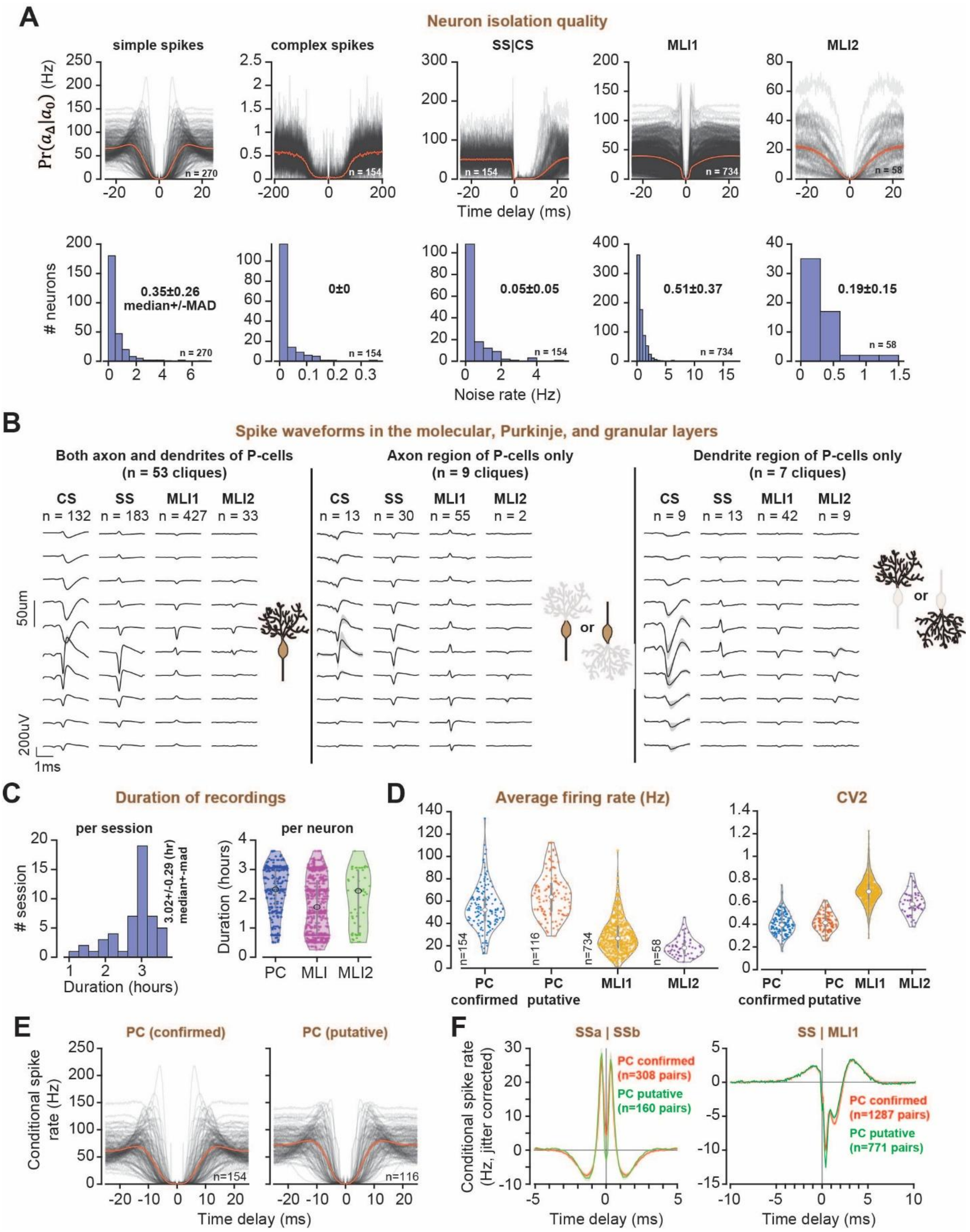


**Extended Data Fig. 1. Isolation quality and duration of recording of the neurons. (A)** Top: High resolution auto-correlograms (and SS|CS cross correlogram) of each cell type with average traces (red). SS|CS describes the conditional probability of an SS, given that a CS occurred at time zero. The probabilities are for bins of 0.1 ms duration and are multiplied by 10000 to represent spike rate in Hz (except for CS|CS which bin size is 2ms). Bottom: refractory violation distribution of for each cell type (<1ms for all neurons except complex spikes and SS|CS: <5ms and CS|CS: <10ms). **(B)** Normalized spatiotemporal aligned waveforms of each cell type (see Extended Data Fig. 2 for more details) plotted as a function of distance of the electrode with respect to the P-cell layer or electrode with maximum CS waveform. The cliques are divided into (left) full P-cell waveforms with both dendritic and somatic spike shape, (middle) cliques with only axonal complex spike waveforms, (right) cliques with only dendritic complex spike waveforms. **(C)** Duration of the neurophysiological recording (50 sessions), and duration of isolation for each neuron. Circles are medians, and the lines are median absolute deviation distance (MAD). **(D)** Average firing rates (left) and CV2 (right) of confirmed and putative P-cells, MLI1s, and MLI2s. **(E)** Autocorrelations of confirmed and putative P-cells. The red trace is the average value. **(F)** Left subplot: Jitter corrected conditional probability of spiking in a P-cell, given that another P-cell spiked at time zero. Data are for neuron pairs that are within the same clique. Right subplot: jitter corrected conditional probability of spiking in a P-cell, given that an MLI1 spiked at time zero. Error bars are SEM.

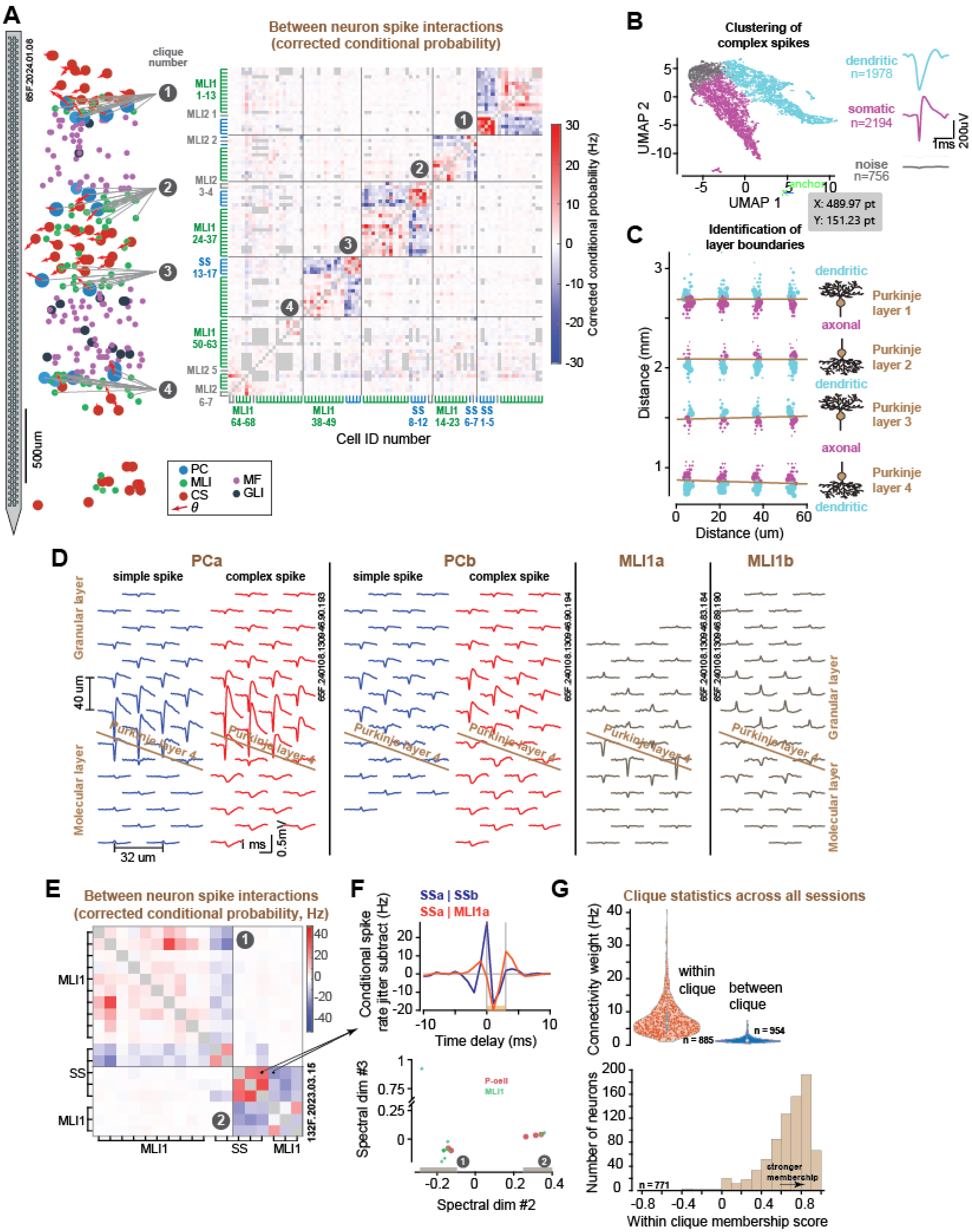


**Extended Data Fig. 2. Organization of neurons into cliques and detection of P-cell layer based on complex spike waveform. (A)** (left) Example Neuropixels session with 4 neuronal cliques. (Right) Adjacency matrix showing simultaneously recorded neurons (rows and columns). Color intensity indicates the strength of jitter-corrected conditional spike interactions. Four cliques were identified. **(B)** Method used for finding the P-cell layer using classification of complex spike waveforms on each electrode into dendritic and somatic spikes using UMAP and Gaussian Mixture Model (waveforms with less than 25uV absolute peak are tagged as noise channels, gray group). **(C)** Complex spike waveforms are color coded by dendritic (cyan) and somatic (magenta) for each electrode recording a complex spike. The size of the points represents the logistic regression weights based on the size of the spike waveform used to find the classification boundary (Purkinje layer). **(D)** Example of finding the P-cell layer. The figure shows the multi-contact waveforms of two example P-cells and pMLI1s from Purkinje layer 4 alongside the estimated Purkinje layer based on complex spike waveform. **(E)** Clique clustering example. Pairwise jitter-corrected cross correlograms were computed for the recorded neurons, resulting in an adjacency matrix. **(F)** Top: Each element of the matrix captures the strength of the interactions between neuron pairs in the 0-3 ms period. Two of the matrix elements are shown here. Bottom: Spectral clustering of the symmetric adjacency matrix separated the two cliques. **(G)** Top: Interaction strength for cell pairs within and between cliques. Bottom: Validation of the clique membership procedure. The plot illustrates the average within versus between clique interaction strength via silhouette-like membership score (see Methods).

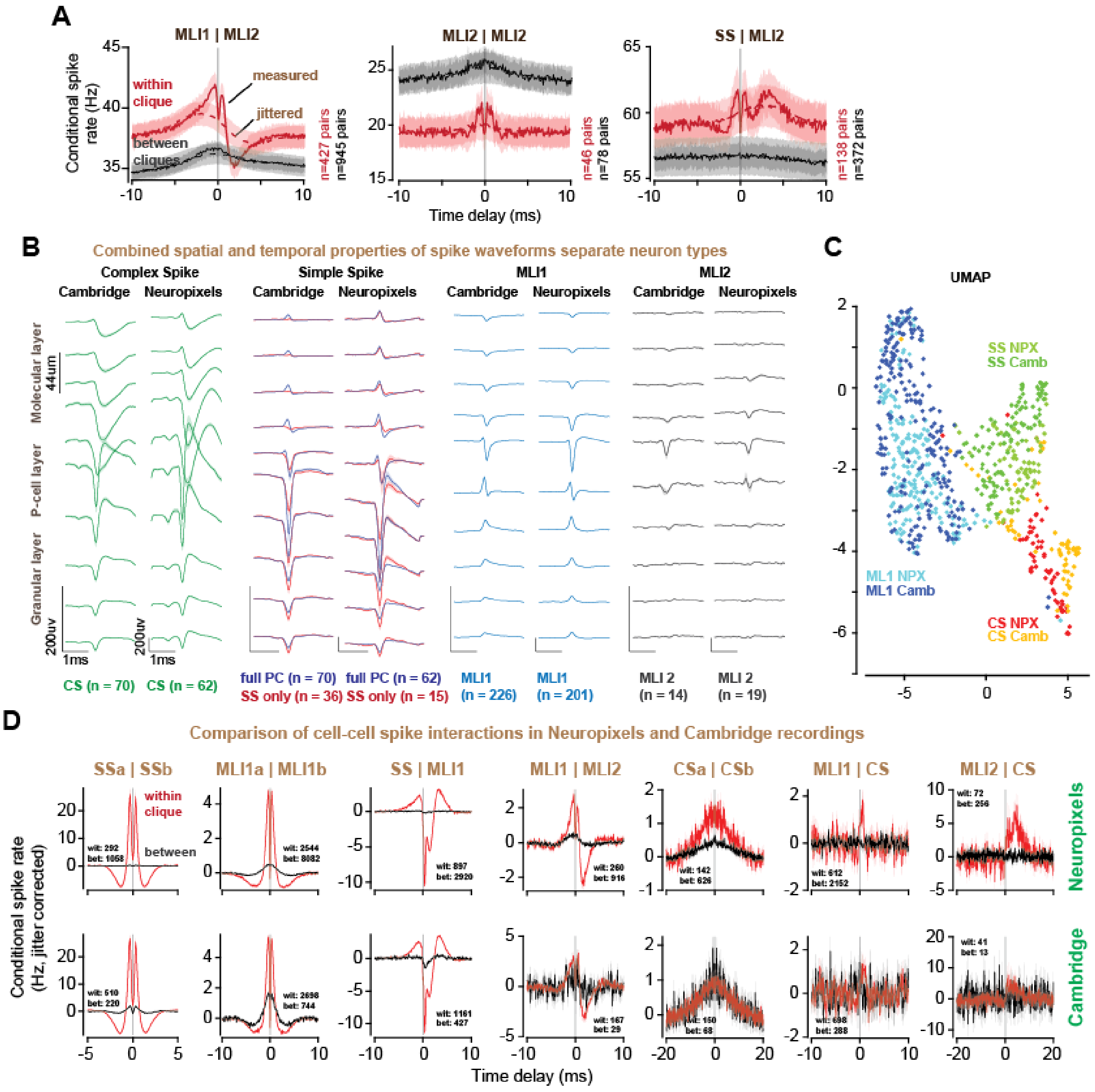


**Extended Data Fig. 3. Interactions of pMLI2 with pMLI1s and P-cells. The spatial patterns of spike waveforms separate neuron types but remain consistent across different silicon probes (Neuropixels and Cambridge). (A)** pMLI1 and pMLI2 interactions: spike probability in pMLI1 given that a pMLI2 spike occurred at 0ms for pairs within and between cliques, alongside jittered traces (dashed lines). Also shown are pMLI2|pMLI2, and SS|pMLI2. **(B)** Waveforms for complex spikes, simple spikes, pMLI1 spikes, and pMLI2 spikes recorded by each probe. The dendritic region is toward the upper part of the figure, the P-cell layer is near the middle, and the granular layer is toward the bottom of the figure. Note the positive spike waveform for pMLI1s near the P-cell layer, consistent with a pinceau. Also note that the voltage scale for Cambridge is nearly 5 times larger than Neuropixels. **(C)** Clustering of the neurons based on their spatiotemporal spike waveforms using UMAP. In these two probes, the complex spikes, simple spikes, and pMLI1 spikes tend to cluster together. **(D)** The spike interactions among neurons remained consistent across different silicon probes.

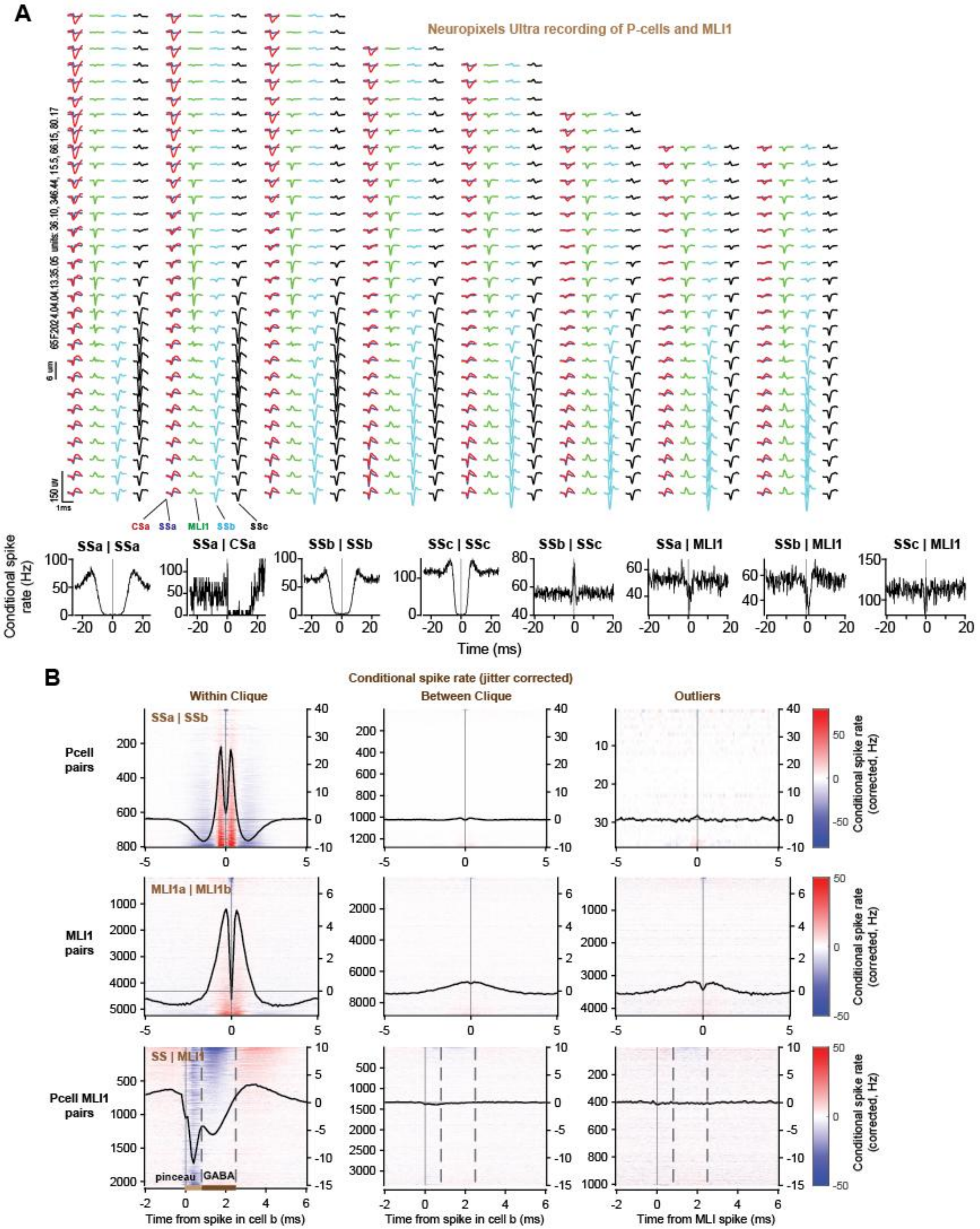


**Extended Data Fig. 4. A Neuropixels Ultra recording of P-cells and MLI1s. (A)** The figure shows 3 P-cells and one pMLI1. One of the P-cells has both simple and complex spikes. The molecular layer is in the upper part of the figure, and the granular layer is in the lower part. Note the positive MLI1 spike suggesting a pinceau. Spike temporal interactions are shown in the lower part of the plot. Data from a 10-minute duration recording. **(B)** High-resolution jitter-corrected conditional probabilities for all cell pairs within-clique, between-clique, and outliers. Heat map and average of jitter-subtracted interactions between P-cell|P-cell (top), pMLI1|pMLI1 (middle), and P-cell|pMLI1 (bottom) pairs within (left), between (middle) cliques and outliers (right). The pairs are sorted by average values within the -1ms to +1ms period for P-cell|P-cell and pMLI1|pMLI1 pairs and 0.8 to 2.5ms (GABA suppression period) for P-cell | pMLI1 pairs.

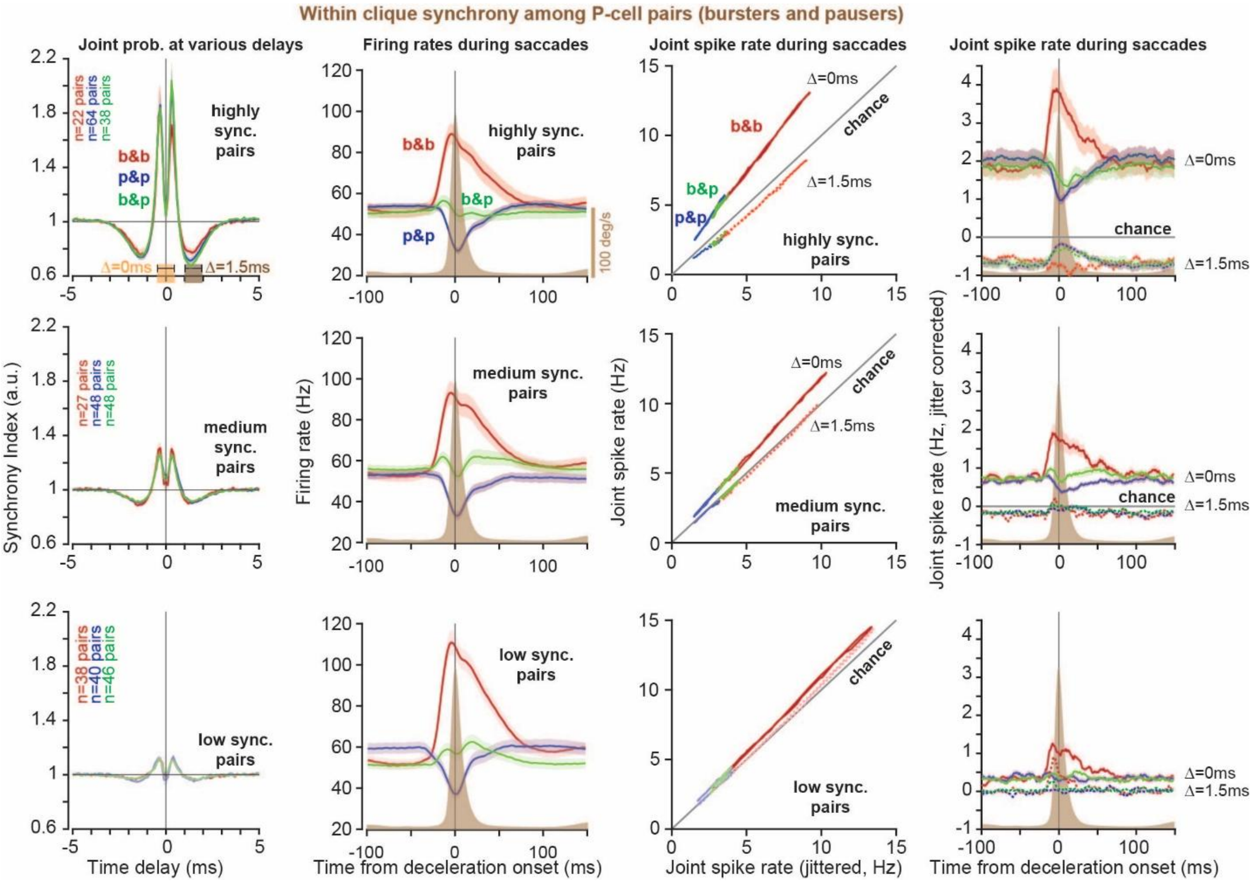


**Extended Data Fig. 5. Joint spike rates at various delays during saccades in pairs of P-cells, organized by combinations of burster/pauser P-cell pairs based on strength of ephaptic coupling.** (Left to right) Jitter corrected whole recording cross-correlograms, average pair firing rates aligned to saccade deceleration onset, joint-jitter plots, and corrected joint probability for burster-burster (red), pauser-pauser (blue), and burster-pauser (green) pairs. Pairs are divided into high, medium, and low synchrony index groups from top to bottom. Note that in the burster pairs with greatest strength of ephaptic coupling (high synch pairs), joint probability of synchronous spikes at $\Delta = 0$ms peaks at saccade deceleration onset while the probability of slightly asynchronous spikes at $\Delta = 1.5$ms is suppressed below chance.

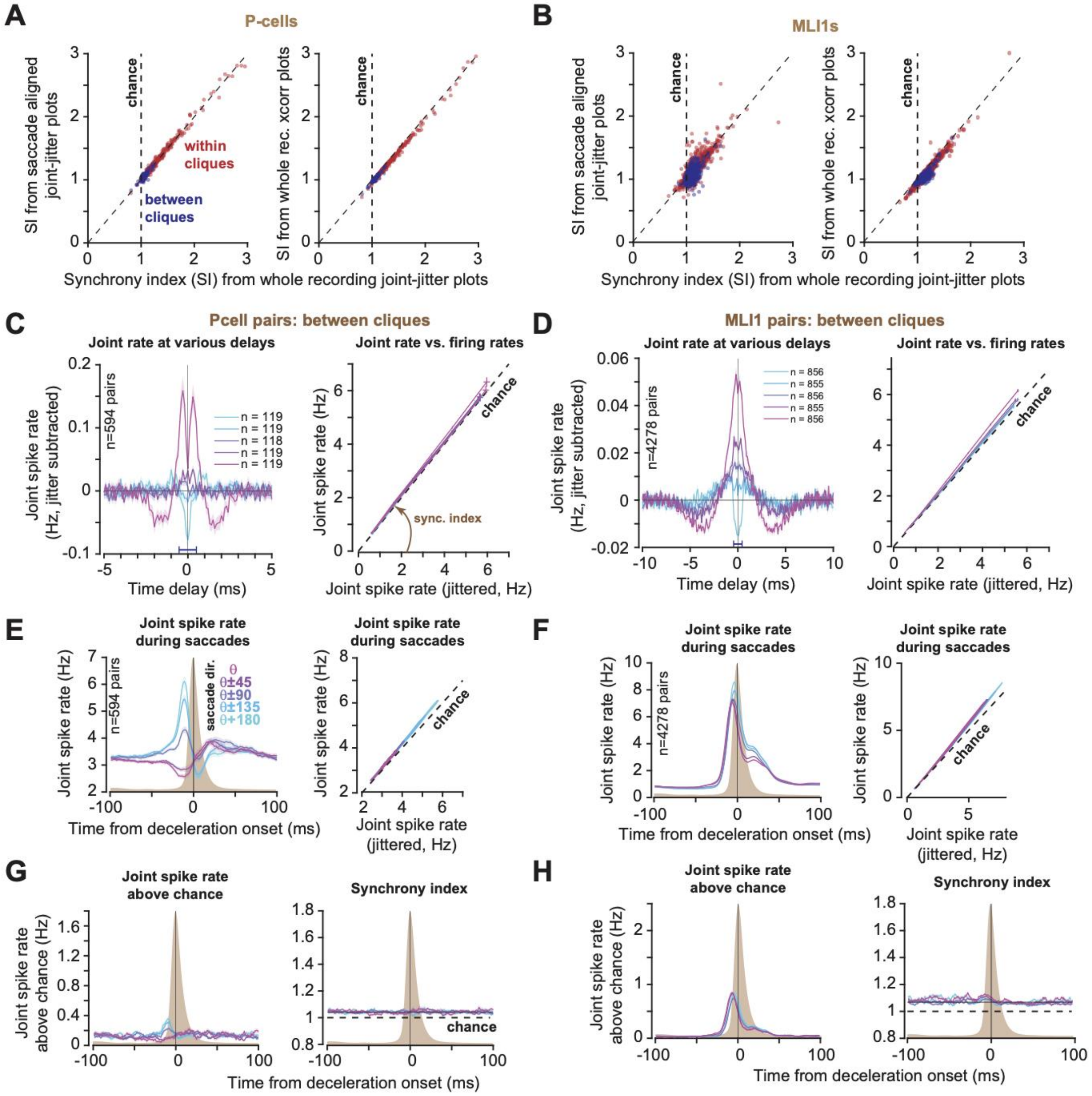


**Extended Data Fig. 6. Three different methods of calculating synchrony index produce similar values and between clique synchrony during behavior and whole recording. (A)** Synchrony index from saccade aligned joint jitter plots (left) and synchrony index from whole recording cross-probabilities (right) versus synchrony index from whole recording joint-jitter plots for P-cells. **(B)** same as (A) for pMLI1 pairs. **(C)** Jitter corrected joint spike rate of P-cell pairs between cliques for different delays (left), and joint-jitter plots (right) binned by synchrony index (compare with Fig. 3 within clique). **(D)** same as (C) for pMLI1 between clique pairs. **(E)** Joint spike rate (left) and joint-jitter plots (right) aligned to saccade deceleration onset for different saccade directions aligned to the potent vector. **(F)** same as (E) for pMLI1 between clique pairs. **(G)** synchronous spike rate above chance (left) and synchrony index aligned to saccade deceleration onset (right). **(H)** same as (G) but for between clique pMLI1s.

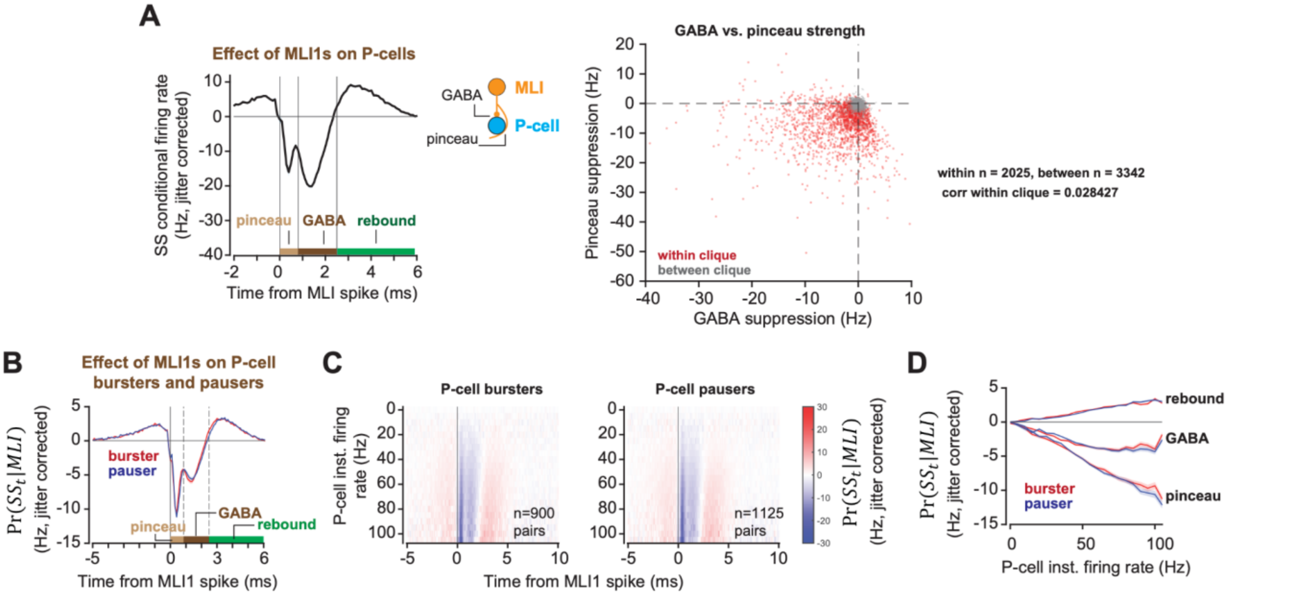


**Extended Data Fig. 7. Independent pMLI1 suppression of P-cells during pinceau and GABA period and similar suppression of P-cell bursters and pausers. (A)** (left) example P-cell conditional firing rate aligned to pMLI1 spike, (right) for each pair of MLI1 and P-cell, this plot displays the strength of pinceau period suppression with GABA period suppression. Pinceau period is 0-0.8ms following MLI1 spike, while the GABA period is 0.8-2.5ms. **(B)** Jitter corrected cross-correlograms between P-cell bursters (red) and pausers (blue) and pMLI1s for the whole recording period. **(C)** Same as (B) but 3D cross-correlograms binned by P-cells instantaneous firing rates. **(D)** Pinceau (0-0.8ms), GABA (0.8-2.5ms), and rebound (2.5-6ms) period average effect for bursters (red) and pausers (blue) as a function of P-cell firing rate.

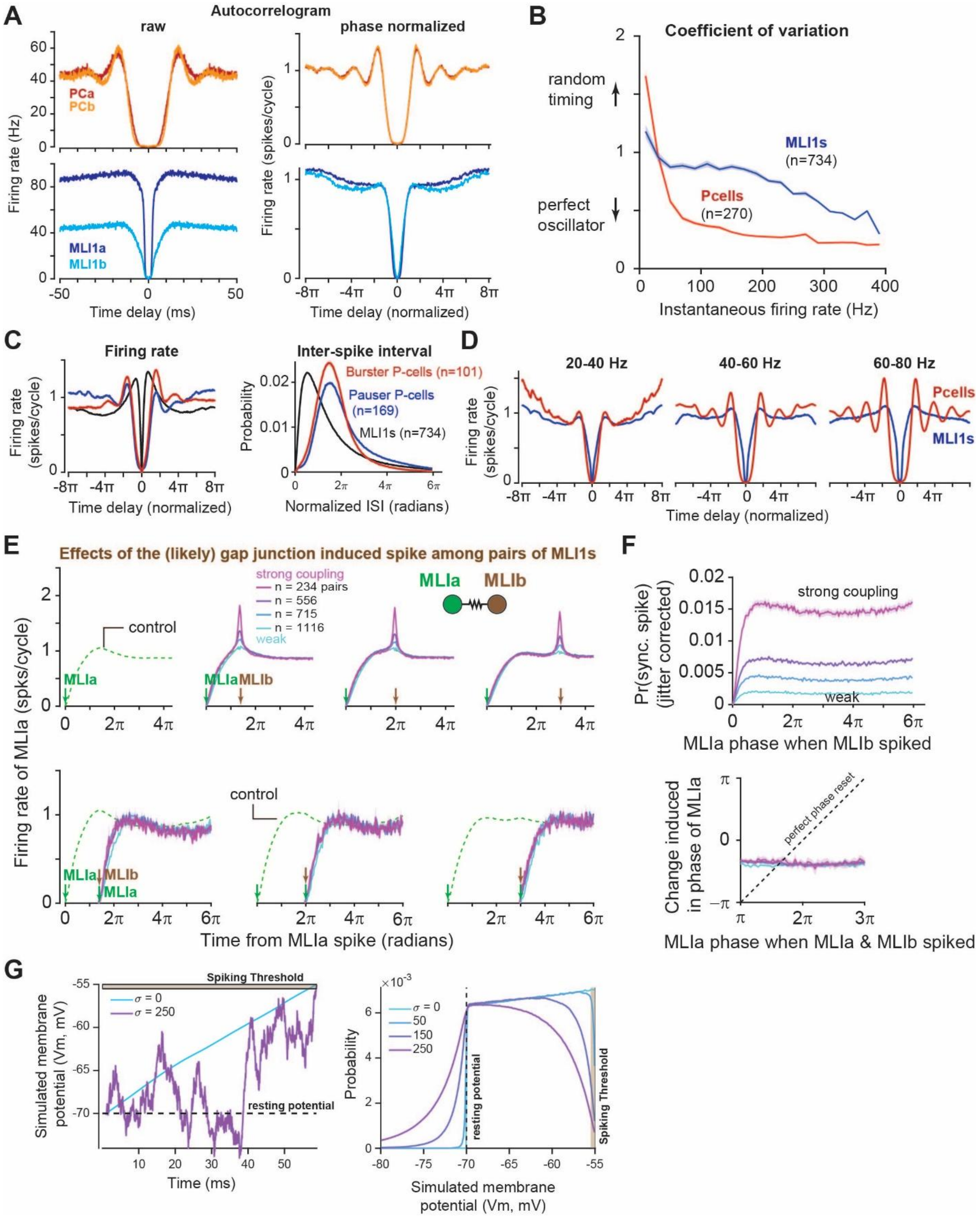


**Extended Data Fig. 8. P-cells show regular spiking patterns unlike pMLI1s. (A)** Auto correlogram (left) and phase-normalized auto correlogram (right) for two example P-cells (top) and pMLI1s (bottom) presented in Fig, 2a-c. **(B)** Coefficient of variation as a function of instantaneous firing rate for all P-cells (red) and pMLI1s (blue). Values are calculated using the 3D ISI distributions in Fig. 5a. **(C)** Average phase-normalized auto correlogram (left) and phase-normalized inter-spike interval distribution (right), using only those spikes that occur within 30 ms of the peak velocity of a saccade. **(D)** Average phase-normalized autocorrelograms for all P-cells at sample instantaneous firing rates near the baseline rate for a typical P-cell. **(E)** Phase resetting control among pairs of pMLI1s. Analogous to Fig. 6c, but for pMLI1s instead of P-cells. (Top) Conditional probability of MLIa firing at each phase given a neighboring MLIb within a clique spiked at a given phase, grouped by different MLIa and MLIb gap junction strengths: from left to right: Control, MLIb at 7π/5, 2π, and 3π. (bottom) Probability of MLIa firing at each phase given MLIb spiked at a given phase $\Delta$ with respect to MLIa and induced a synchronous spike in MLIa: from left to right Δ=7π/5, 2π, and 3π. **(F)** Analogous to Fig 6d. (left) Probability of synchronous spike (jitter corrected joint spiking of MLIa and MLIb) given MLIb spiked at different phases with respect to MLIa, grouped by gap junction strength. MLIa phase has no impact on the chance that an MLIb spike will induce a synchronous spike, except during the refractory period. (right) Expected change in phase of MLIa as a function of the phase when MLIa and MLIb spiked synchronously. The expected change is constant with respect to the phase when the synchronous spike occurred, suggesting that phase carries no information for irregular cells such as MLIs. **(G)** Simulation results: Regularity increases the time spent in the ephaptic spiking band. Left: Example simulated membrane voltage traces for a neuron with no noise (blue) and with high noise (magenta). The gold shaded region is the ephaptic spiking band, the range of voltages within which the small voltage perturbation caused by an ephaptic interaction may be sufficient to induce a spike. Right: Distribution of membrane voltage values for four simulated neurons with different noise levels. The frequency of occurrence of membrane voltage values near threshold decreases with increasing noise. Error bars are SEM for all figures, except F bottom, where the shaded region is the 95% confidence interval on the circular mean, obtained by bootstrapping on the population of pMLI1 pairs.

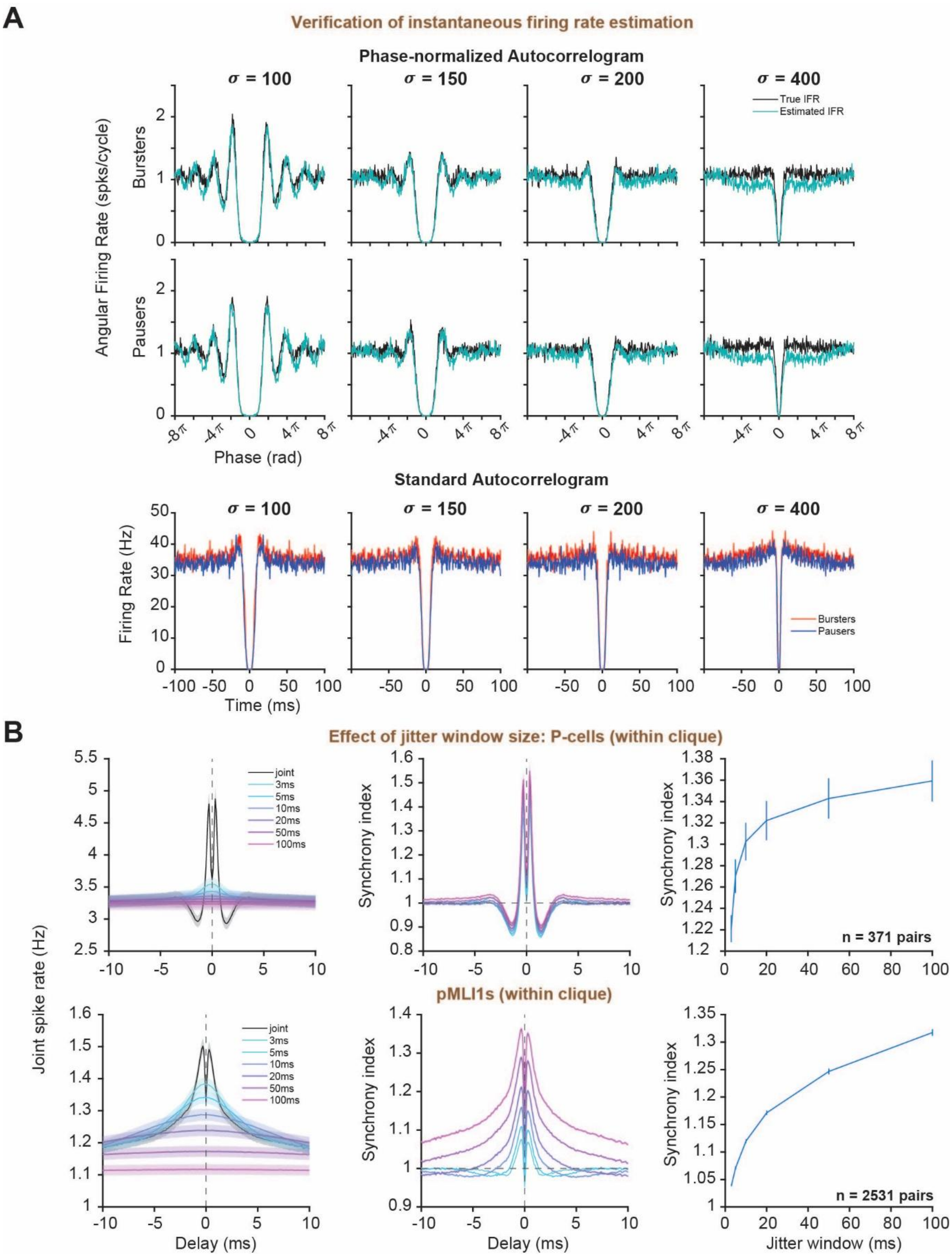


**Extended Data Fig. 9. Validation of hyperparameter selections for instantaneous firing rate phase normalization methods and jittering window. (A)** Top: Conditional spiking probability as a function of phase for simulated spike trains with various levels of noise: left to right: σ = 100, 150, 200, 400. The spike trains were generated with target firing rates given by concatenation of the trial-averaged firing rates for each recorded burster P-cell, aligned to the time at which the eye reached maximum velocity for each saccade. Phase was measured using two different firing rates: the target firing rates that were used to generate the spike trains (black), and the estimated instantaneous firing rate found by smoothing the inverse of the inter-spike intervals, as was used to measure phase for the real data (blue). The results show minimal bias in using the estimated instantaneous firing rate compared to the ground truth firing rate. Middle: Same analysis applied to simulated spike trains that were generated by concatenation of trial-averaged pauser firing rates. Bottom: Standard auto correlograms for the simulated burster and pauser spike trains. These figures demonstrate how the regularity that is apparent in the phase-normalized conditional firing plots can be obscured due to variations in firing rate over time, explaining why some regular real P-cells do not show obvious signs of oscillatory behavior in their auto correlograms. **(B)** Effect of jittering window on measured synchrony values for within clique P-cell (top row) and pMLI1 (bottom row) pairs. (left) Average joint firing rate of two P-cells for different jittering window values, (middle) same as (left) but normalized by jittered rates (synchrony index), (right) synchrony index values (average from -0.5 to 0.5ms delay) for each jittering window size for pairs within cliques.

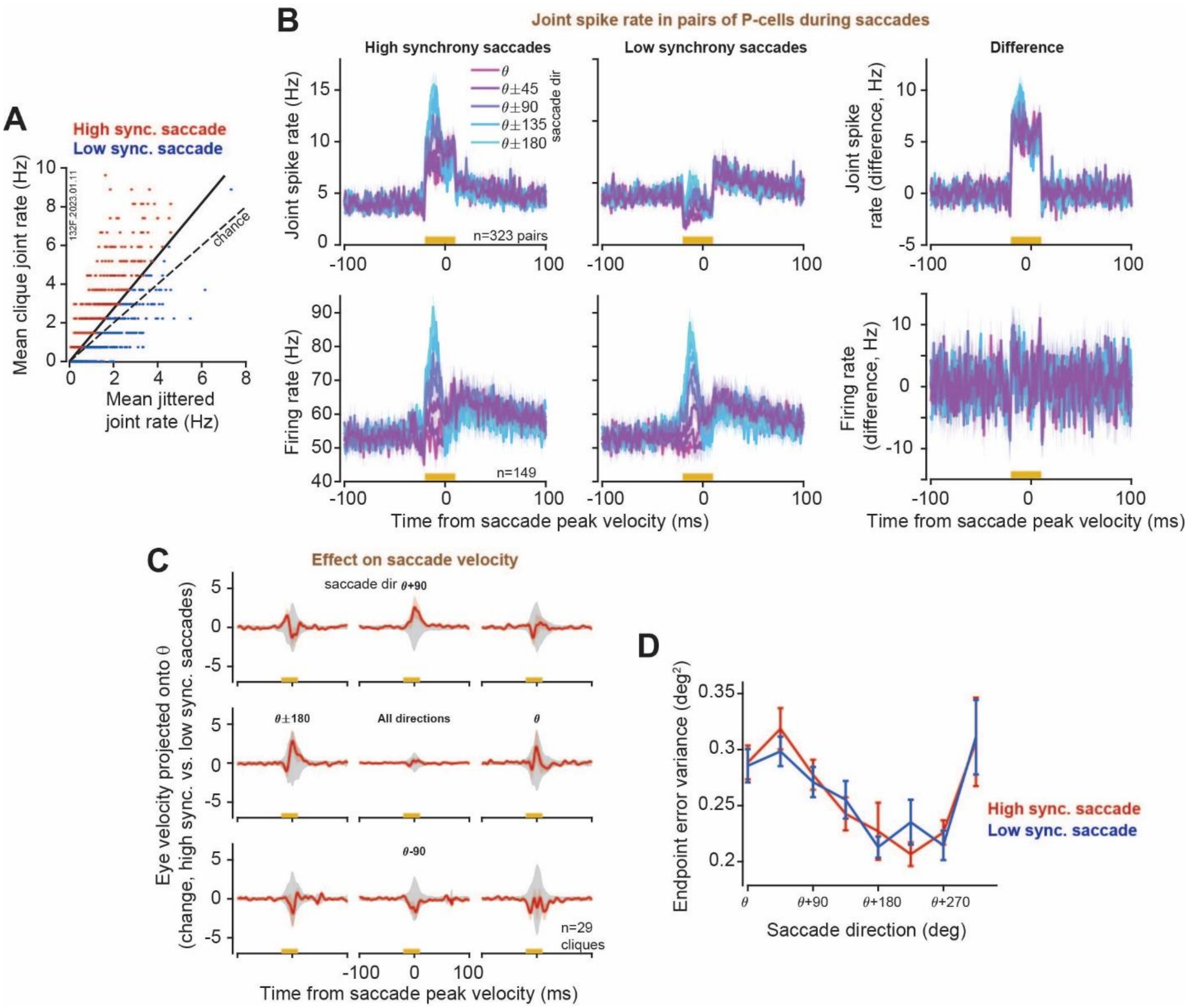


**Extended Data Fig. 10. Presence of synchronous spikes in single pairs of P-cells did not produce measurable effects on saccades**. **(A)** Mean jittered joint rate vs true mean joint rate for individual trials, averaged across all P-cell pairs in an example clique. Trials are classified as high synchrony (red) or low synchrony (blue). The slope of the solid line indicates the mean synchrony index of the pairs in the clique. **(B)** Top: Average synchronous P-cell firing rate for high synchrony trials (left), low synchrony trials (middle), the difference between high and low synchrony (right). The gold bar indicates the window that was used to classify trials. Bottom: Individual P-cell firing rates for the same trial conditions. **(C)** Mean eye velocity difference, high synchrony trials – low synchrony trials, for saccades in eight different directions with respect to the clique CS-on direction ($\theta$). Gray shaded region indicates 95% confidence interval, obtained by shuffling the labels of high and low synchrony trials. Results are averaged across all cliques that contain at least 4 P-cells and weighted by the number of P-cells in each clique. On average, across the eight directions, the difference in eye velocity between high and low synchrony trials was not significant (permutation test, center plot). (D) Variance in endpoint error, averaged across cliques, for high synchrony trials (red) and low synchrony trials (blue). High and low synchrony trials were not significantly different for any of the directions (paired t-test, $p > 0.2$ for all directions). Error bars are SEM.

**Supplementary Table 1. Parameter values for P-cell ephaptic coupling simulations**

| **Parameter** | **Value** |
|---|---|
| **P-cell intrinsic parameters** | |
| Membrane capacitance $(C_m)$ | 20 nF |
| Leak conductance $(g_m)$ | 0.5 $\mu S$ |
| Membrane time constant $(\tau)$ | 40 ms |
| Leak reversal potential $(E_L)$ | -70 mV |
| Refractory period $(\tau_{ref})$ | 1 ms |
| Spike threshold $(V_{th})$ | -55 mV |
| Intracellular spike max amplitude | 35 mV |
| Extracellular spike max amplitude | -0.97 mV |
| Spike shape $\gamma$ kernel parameters | $\alpha = 4, \theta = 0.4$ |
| **P-cell input parameters** | |
| Linear rate-to-current gain $(g_{max})$ | 0.4083 nA/Hz |
| External current standard deviation $(I_\sigma)$ | {50, 150, 250} nA |
| **Ephaptic coupling parameters** | |
| AIS-AIS distance $(d)$ | {2.5, 5, 10, 25, 50} $\mu m$ |
| Geometry/offset constant $(k_0)$ | 320 $\mu m$ |
| **Trial parameters** | |
| Simulation time step | 0.0333 ms |
| Simulation duration | 240 seconds |
| Number of trials per simulation | 500 trials |
| Number of simulations per condition | 120 |

**Supplementary Table 2. Parameter values for MLI gap-junction simulations**

| Parameter | Value |
|---|---|
| **MLI intrinsic parameters** | |
| Membrane capacitance ($C_m$) | 10 pF |
| Leak conductance ($g_m$) | 2.5 n$S$ |
| Membrane time constant ($\tau$) | 4 ms |
| Leak reversal potential ($E_L$) | -70 mV |
| Refractory period ($\tau_{ref}$) | 0.5 ms |
| Spike threshold ($V_{th}$) | -55 mV |
| **MLI input parameters** | |
| AMPA $\tau_1$ | 0.1659 ms |
| AMPA $\tau_2$ | 2.9381 ms |
| Glutamatergic reversal potential ($E_{ex}$) | 0 mV |
| Glutamatergic conductance ($g_{e_{max}}$) | 0.25 n$S$ |
| External current mean ($I_c$) | 0.028 nA |
| External current standard deviation ($I_\sigma$) | 0.016 nA |
| Gap junction conductance ($g_{gap}$) | {0, 1, 2.5, 5} nS |
| **PF parameters** | |
| Refractory period ($\tau_{ref}$) | 0.5 ms |
| Num. MF inputs per PF | 3-5 neurons |
| Num. PF | 10 neurons |
| **Trial parameters** | |
| Simulation time step (dt) | 0.0333 ms |
| Simulation duration | 500 seconds |
| Num. behavioral trials | 3 saccades per second |
| Num. simulations per condition | 120 |
